# Tissue confinement regulates cell growth and size in epithelia

**DOI:** 10.1101/2022.07.04.498508

**Authors:** John Devany, Martin J Falk, Liam J Holt, Arvind Murugan, Margaret L Gardel

## Abstract

Cell proliferation is a central process in tissue development, homeostasis and disease. Yet how proliferation is regulated in the tissue context remains poorly understood. Here, we introduce a quantitative framework to elucidate how tissue growth dynamics regulate cell proliferation. We show that tissue growth causes confinement that suppresses cell growth; however, this confinement does not directly affect the cell cycle. This leads to uncoupling between rates of cell growth and division in epithelia and, thereby, reduces cell size. Division becomes arrested at a minimal cell size, which is consistent across diverse epithelia *in vivo*. Here, the nucleus approaches a volume limit set by the compacted genome. The loss of Cyclin D1-dependent cell size regulation results in an abnormally high nuclear-to-cytoplasmic volume ratio and DNA damage. Overall, we demonstrate how epithelial proliferation is regulated by the interplay between tissue confinement and cell size regulation.

**Highlights:**

- In epithelia, regulation of cell growth and cycle are uncoupled
- Cell growth is regulated by tissue-scale dynamics, which determine confinement
- Cell volume in epithelial tissue is described by G1 sizer model with a tunable growth rate
- Volume of cells in epithelial tissues is near a minimum set by genome size

## INTRODUCTION

Regulation of cell proliferation is a central question for understanding tissue development, growth and homeostasis (Murray and Hunt, 1993; Nurse, 2000). In contrast to the exponential proliferation of isolated cells (Broach, 2012; Scott and Hwa, 2011), multicellular tissue requires tight coupling between cell proliferation and tissue growth (Irvine and Shraiman, 2017). One proposed mechanism for regulating proliferation in epithelia is the so-called process of “contact inhibition of proliferation” (henceforth referred to as contact inhibition) where cell proliferation becomes highly restricted due to spatial constraints imposed by the tissue (Irvine and Shraiman, 2017; McClatchey and Yap, 2012). The disruption of contact inhibition results in cell overgrowth and altered tissue architecture (Fomicheva and Macara, 2020; Kim et al., 2011; Leontieva et al., 2014). Therefore, contact inhibition is thought to play a key role in maintaining tissue homeostasis and preventing tumor formation (Irvine and Shraiman, 2017; Mendonsa et al., 2018). However, because contact inhibition is regulated through multiple signaling pathways and largely unknown parameters, we currently lack a framework for understanding the process across diverse tissues. This is evident from the literature where contact inhibition is described as dependent on cell density (Ibar et al., 2018), adhesion signaling (Kim et al., 2011; McClatchey and Yap, 2012) and mechanical stress (Irvine and Shraiman, 2017; Pan et al., 2016). As these variables are difficult to manipulate independently, it remains unclear how tissue geometry and growth dynamics impacts size and growth of constituent cells.

The regulation of size and growth of isolated mammalian cells is, by contrast, well understood (Cadart et al., 2018; Tan et al., 2021; Zatulovskiy et al., 2020; Zatulovskiy and Skotheim, 2020). Single cells go through cycles of coupled growth and division and have stable cell size due to feedback between cell growth and division. However, there appear to be different feedback mechanisms acting in different contexts. For example, single cells mainly sense the total amount of cell growth throughout the cell cycle to regulate the length of cell cycle phases (Cadart et al., 2018; Tan et al., 2021). In contrast, mouse epidermal tissue shows feedback via a cell size checkpoint preventing small cells from entering S phase (Xie and Skotheim, 2020). Due to limited data *in vivo* it remains unclear if this is a consequence of tissue specific behavior in the skin or could reflect a difference in regulation between single cells and epithelial tissue. Several lines of evidence suggest the latter is true. For instance, the cell cycle duration can be on the scale of weeks or months *in vivo* (Sender and Milo, 2021), challenging established dilution-based mechanisms of cell cycle regulation (Zatulovskiy et al., 2020; Zatulovskiy and Skotheim, 2020). Further, prior work has shown a possible switch to size-dependent regulation of the cell cycle in cell culture models of epithelial tissue (Puliafito et al., 2017, 2012). However, we lack a systematic study that explores how tissue-scale growth dynamics impact regulation of cell size and growth in epithelia.

To understand how cell proliferation is regulated in epithelium, we first performed a meta-analysis of cell size data and found that epithelial cell sizes are remarkably consistent and highly context dependent. The cell size in epithelial tissue *in vivo* is always smaller than single cells in culture. However, we could recapitulate the *in vivo* cell size using a cell culture model of epithelial tissue formation, indicating that tissue-scale phenomena impact cell size regulation. To quantitatively explore this, we employed model epithelial tissues with varied growth rates. We then introduce a general framework to quantify how tissue-scale growth dynamics constrain cell growth, providing a measure of “tissue confinement”. We show that increasing tissue confinement reduces cell growth and YAP/TAZ signaling but does not impact the cell cycle duration directly. Instead, cell cycle duration is regulated by cell size. There is a sharp cell cycle arrest at a cell volume of ∼1000 µm^3^, consistent with cell size found *in vivo*. In both epithelial and single cell contexts, cell size regulation is well described by a “G1 sizer model” with a tunable growth rate. In epithelia cyclin D1 protein levels are strongly cell size-dependent and overexpression of cyclin D1 reduces the minimal cell size. This suggests that the levels of cyclin D1 controls size-dependent cell cycle arrest. We see that abnormally small cells display DNA damage as these cells approach a size limit set by the volume occupied by the fully compacted genome. This suggests that, in addition to mediating cell cycle arrest during contact inhibition, cell size regulation pathways are critical for maintaining epithelial homeostasis. Overall, we demonstrate the general mechanisms of cell growth and division regulation in epithelia which provides new insight into proliferative processes in tissue development, homeostasis, and disease.

## RESULTS

### Epithelial cell size is context dependent

To screen for varied cell size regulation in epithelial tissue, we systematically compared cell volume from different tissues *in vivo* and cell culture models (Table S1). We compiled these data from available sources including histology sections from the human protein atlas (Fig. 1A) (Uhlén et al., 2015), 3D segmentation data from Allen Cell Institute (Fig. 1A) (3D cell viewer - Allen Cell Explorer.) and published cell volume measurements (Cadart et al., 2018; Elamin et al., 2012; Engelberg et al., 2011; Goldspink et al., 2017; Guo et al., 2017; Karve et al., 2017; Leung et al., 2014; Medeiros et al., 2021; Morizane et al., 2015; Padovan-Merhar et al., 2015; Park et al., 2010; Perez-Gonzalez et al., 2019; Schutgens et al., 2019; Viana et al., 2021; Yao et al., 2020; Zhao et al., 2008). When cell volume measurements weren’t available, we estimated cell volume by identifying cells perpendicular to the tissue slice and then measuring their length and width. We confirmed that this provides an accurate estimation of volume from benchmarking against full 3D imaging (Fig. S1&S2). Across 15 tissue types in the human protein atlas, we find the volume of epithelial cells is surprisingly consistent, with a narrow distribution of 630±180 μm^3^ (Fig. 1B, circles). By comparison, the volume across 12 types of isolated epithelial or epithelial-like cells is consistently measured to be severalfold larger, with a volume of 2330±650 μm^3^ (Fig. 1B, squares) (Cadart et al., 2018; Guo et al., 2017; Leung et al., 2014; Park et al., 2010; Perez-Gonzalez et al., 2019; Viana et al., 2021; Zhao et al., 2008). Interestingly, the volumes of epithelial cells cultured in 3D more closely matched those *in vivo*, with a mean of 1020±520 μm^3^ (Fig. 1B, diamonds) (Elamin et al., 2012; Engelberg et al., 2011; Goldspink et al., 2017; Karve et al., 2017; Medeiros et al., 2021; Morizane et al., 2015; Schutgens et al., 2019). These data suggest that epithelial cell volume is strongly influenced by the tissue environment.

**Figure 1:**
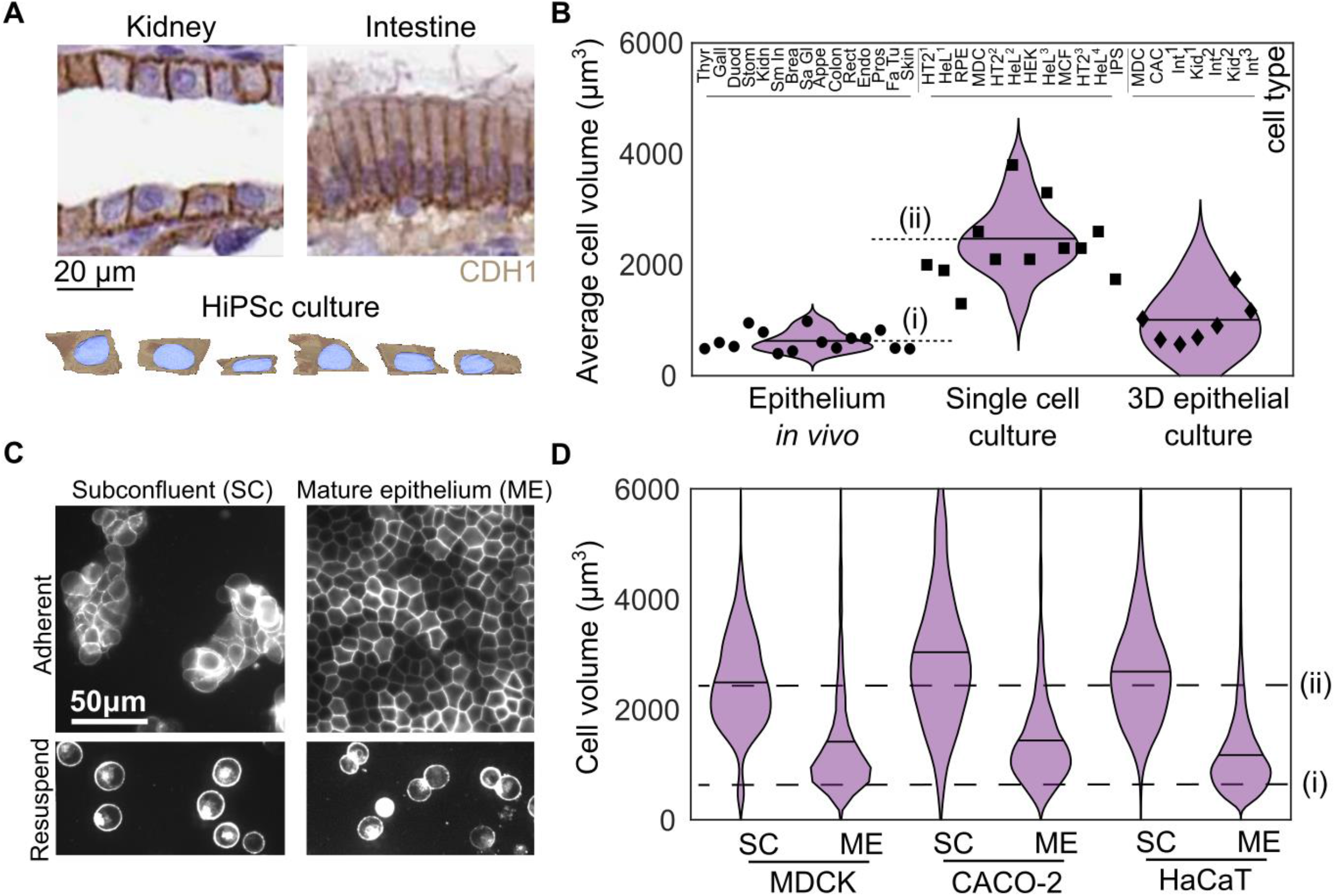
Epithelial cell size is consistent across tissues but context dependent. (A) Histology sections published by the Human Protein atlas showing E-cadherin (CDH1) staining kidney and Intestine (Duodenum) tissue (image credit: Human Protein Atlas) and human induced pluripotent stem cell (HiPSc) segmentation data from Allen Cell Institute (Available from allencell.org/3d-cell-viewer). (B) Cell volume from 15 tissues in vivo (circles), 12 published measurements from cell lines cultured as single cells (squares) and 7 measurements from 3D epithelial cell cultures (diamonds). Each data point is the average cell size from >50 cells for a given cell type. Measurement details available in Table S1. Dashed lines show the average cell volumes from epithelium in vivo (i) and in single cell culture (ii) (C) (Top) Images of MDCK cells with fluorescently labeled cell membranes (stargazin-gfp) in subconfluent colonies (SC) and mature epithelium (ME). (bottom) Cells under each condition are also shown after being treated with trypsin for 10 minutes and resuspended. (D) Cell volume in SC and ME for MDCK, CACO-2 and HaCaT cell lines. Dashed lines show the average cell volumes from epithelium in vivo (i) and in single cell culture (ii) from B (N = cells (experiments), N_MDCK-SC_ = 5884(4), N_MDCK-ME_ =13513(5), N_CACO2-SC_ = 732(1), N_CACO2-ME_ = 2735(2), N_HaCaT-SC_ =9248(2), N_HaCaT-ME_ =13451(2))

To test this hypothesis, we utilized several different epithelial cell lines (Madin-Darby canine kidney (MDCK), Caco-2 and HaCaT) to measure the cell volume in either subconfluent colonies (SC) or mature epithelium (ME). Controlling cell plating density allowed for the formation of sub-confluent colonies each comprised of 50-1000 cells or a nearly confluent monolayer on collagen gels (see Methods). After the initial plating of monolayers, cell dynamics drive changes in their density, shape and speed over the next 1-2 days (Devany et al., 2021); we define a mature epithelium as the time at which these properties stop changing in time (see Methods). Cell sizes were visualized with a fluorescently tagged membrane protein (stargazin-gfp (CACNG2-gfp) or stargazin-halotag) (Fig. 1C). To facilitate measurement of cell volume, cells were trypsinized, resuspended and then imaged (Fig. 1C, Fig. S1, see Methods). When in sub-confluent colonies, the cell volume of all three epithelial cell types is ∼2800 μm^3^ (Fig. 1D, SC), consistent with the mean value of single cells in Fig. 1B (Fig. 1D, dashed line (ii)). Further, we found that ME culture conditions reduced the cell volume by 60% to plateau at ∼1000 μm^3^ (Fig. S3), a size consistent with *in vivo* data reported in Fig. 1B (Fig. 1D, dashed line (i),). We performed the same experiment on two cell lines which do not form coherent colonies, retinal pigment epithelial cells (RPE-1) and Mouse embryonic fibroblasts (MEF) and did not see size reduction (Fig. S4). This suggests this context-dependency of cell size may be specific to epithelium. Together, these data suggest that contact inhibition qualitatively changes cell size regulation across diverse epithelia.

### Cell size reduction involves an uncoupling of growth and the cell cycle

To query how cell growth and division rates change during the transition from subconfluent colonies to mature epithelium, live cell imaging was used to monitor changes in cell size and number. The data was aligned such that t=0 h denotes the onset of confluence (OC). For all earlier times, cells are in subconfluent colonies. By t > 12 h the cell movement has ceased, and density is constant; we denote this as a mature epithelium (Fig. 2A). We first consider cell division and growth rates for t << 0 h (SC), t ∼ 0 h (OC) and t >> 0 h (ME). Cell division rate, obtained by cell counting measurements, was completely arrested in the ME but only suppressed by ∼40% at OC, as compared to SC (Fig. 2B, black bars). By contrast, the cell growth, obtained by quantifying the rate of protein dilution using pulse chase labeling (see Methods), was suppressed almost entirely at OC (Fig. 2B, purple bars). These data indicate suppression of cell growth and cycle are not tightly coupled during the transition from subconfluent to confluent tissue. Instead, cell growth is suppressed acutely at OC whereas cell cycle progression is only impacted in later stages (Fig. 2C). In single cells, cell growth and division are coupled to maintain a constant size (Zatulovskiy and Skotheim, 2020). A division rate exceeding the growth rate would, instead, be expected to result in cell size reduction.

**Figure 2:**
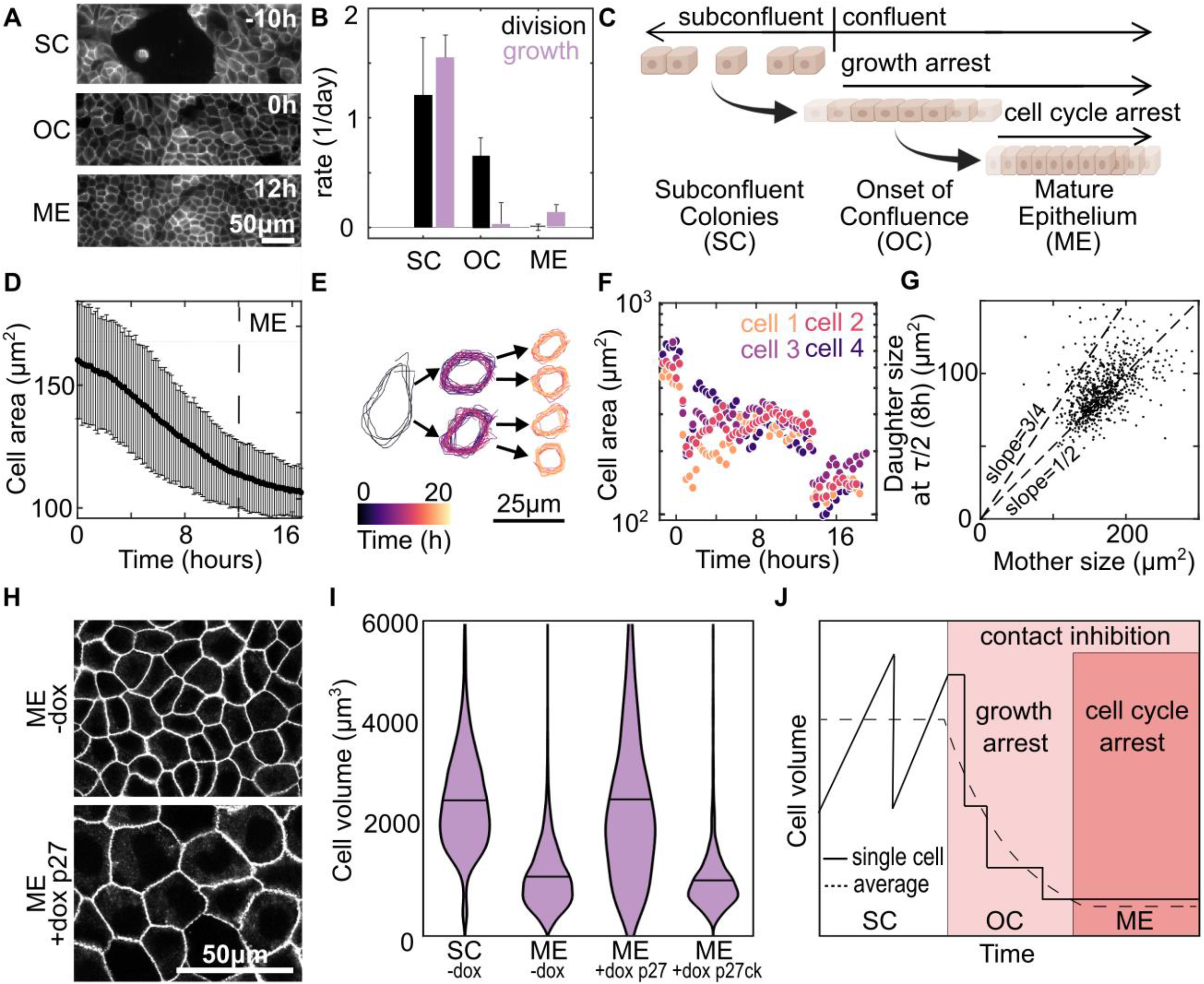
Uncoupling of division and growth at the onset of confluence leads to cell size reduction. (A) Images of MDCK cell membranes at time points during the transition from subconfluent colonies (SC) (t=-10h) to mature epithelium (ME) (t = 12h). The data are aligned such that onset of confluency (OC) occurs at t=0 h. (B) Average rates of cell division (black) and growth (purple) at SC,OC and ME. For growth data protein dilution is measured by CellTrace, n= 10 fields of view with >100 cells from 1 experiment; Error bars are S.E.M. of the fields of view. For division rate, the change in cell number, determined by a cell counter, are averaged from 3 experiments. Error bars are S.D. of experiments (C) Schematic summarizing result that, at the onset of confluence, there is a temporal decoupling of growth and cell cycle arrest. (D) Average cell area over time, t=0 is OC. Error bars are S.D. of 30 fields of view each containing >500 cells; Time at which ME forms indicated by dashed line. (E) Outline of a representative cell, and subsequent daughter cells, over the experiment. (F) Areas of 4 representative cells over the course of the experiment. Traces are shifted in time so that the first cell division occurs at time 0 for all cells. (G) Area of daughter cell halfway through the cell cycle versus the area of mother cell (N= 758 cells from 1 experiment. Linear fit slope = 0.52) (H) Images of cell membrane at t = ME+3 days in monolayers formed from Tet-On P27 cells. ME were formed under control conditions (-dox) or with the cell cycle inhibited (+dox p27) by the addition of doxycycline at t=0 h (I) Cell volumes in SC and ME with Tet-On p27 or Tet-On p27ck (non-cell cycle inhibiting control) in control conditions (-dox) or in ME with doxycycline added at t=0hr. SC and ME -dox data also displayed in Fig. 1D (N_SC_=5884(4) N_ME_=13513(5) N_ME+p27_= 1606(3) N_ME+p27ck_= 1930(3)). (J) schematic of the mechanism of cell volume regulation during the transition from SC to ME

To test this hypothesis, we measured the cell size between t=0 and 18 h during the transition from OC to ME. Over this time, the cell height remained constant, such that changes in cell area were accurate indicators of cell volume (Fig. S2C). The average cell area from a population of ∼1000 cells decreases by ∼50% from OC (t = 0 h) to ME (Fig. 2D). To determine the mechanism driving changes in cell volume we performed single cell tracking. Cells that divided near the beginning of the experiment (t=0 h) were identified and tracked through the experiment. Individual cell trajectories revealed that the cell size reduction occurred by successive cell division events in the absence of cell growth (Fig. 2E, F). Indeed, across a large population of mother-daughter cell pairs, the cell size of a daughter cell 8 hr (∼1/2 cell cycle time, τ) post cell division remained approximately half that of the mother cell size (Fig. 2G). This indicates minimal cell growth during the cell cycle. This is in stark contrast with subconfluent cells, where a cell grows at a constant rate and doubles in size prior to division into two daughter cells (Cadart et al., 2018). For single cells, we would expect the cell to grow by ∼50% at 8 hours post division, resulting in a slope of ∼3/4 (Fig. 2G, dashed line).

To demonstrate that cell division is necessary for cell size to decrease, experiments similar to those in Fig. 2A were performed with a Tet-On p27 MDCK cell line to artificially arrest the cell cycle in the presence of doxycycline (Sherr and Roberts, 1999). In the absence of dox (-dox), the volume decreases in ME compared to SC, similar to our previous experiments (Fig. 2H, I). Addition of doxycycline at OC prevented cell volume decrease in ME (ME +dox p27), and the volume of these cells remained similar to that of SC (Fig. 2H, I). Induction of a mutant p27 which does not arrest the cell cycle (+dox p27ck) (Vlach, 1997) had no effect on cell size in ME (Fig. 2I). Together, these data demonstrate that temporal uncoupling of cell growth and cell cycle result the cell size reduction during formation of mature epithelial tissue. At the onset of confluence, cell growth is highly suppressed and cell volume reduces by division in absence of cell growth (Fig 2J). Then at later times the cell cycle also becomes arrested, and cells reach a final cell size which is comparable to epithelium *in vivo*. We next sought to understand regulation of cell growth and cycle in the epithelium.

### Tissue confinement regulates cell growth

These data motivate the need to introduce a quantitative framework to relate the tissue and cell growth rates. We first consider the growth of a multi-cellular tissue with initial area *A*_0_ and time-dependent area, *A*(*t*). Then, consider this tissue broken up into individual cells to represent the proliferation dynamics of this tissue in the absence of spatial constraints (Fig. 3A). The collection of single cells grows in total size exponentially, such that the total time-dependent area is described by *A*_*U*_(*t*) ∼ 2^*t*/*τ*^, where *τ* is the average cell cycle time. The deviation of the tissue growth from this hypothetical maximum exponential rate quantifies the constraints tissue growth places on cell growth. For a certain tissue area, *A’*, if the ratio of the tissue growth rate to the unconfined growth rate: 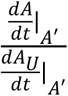 is less than 1, then the tissue growth dynamics constrain cell proliferation. We then define the tissue confinement 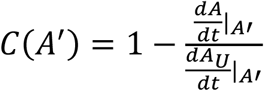 such that there is no confinement effect (C=0) when single cells and tissues have identical growth dynamics and C=1 when tissue growth rate is zero.

**Figure 3:**
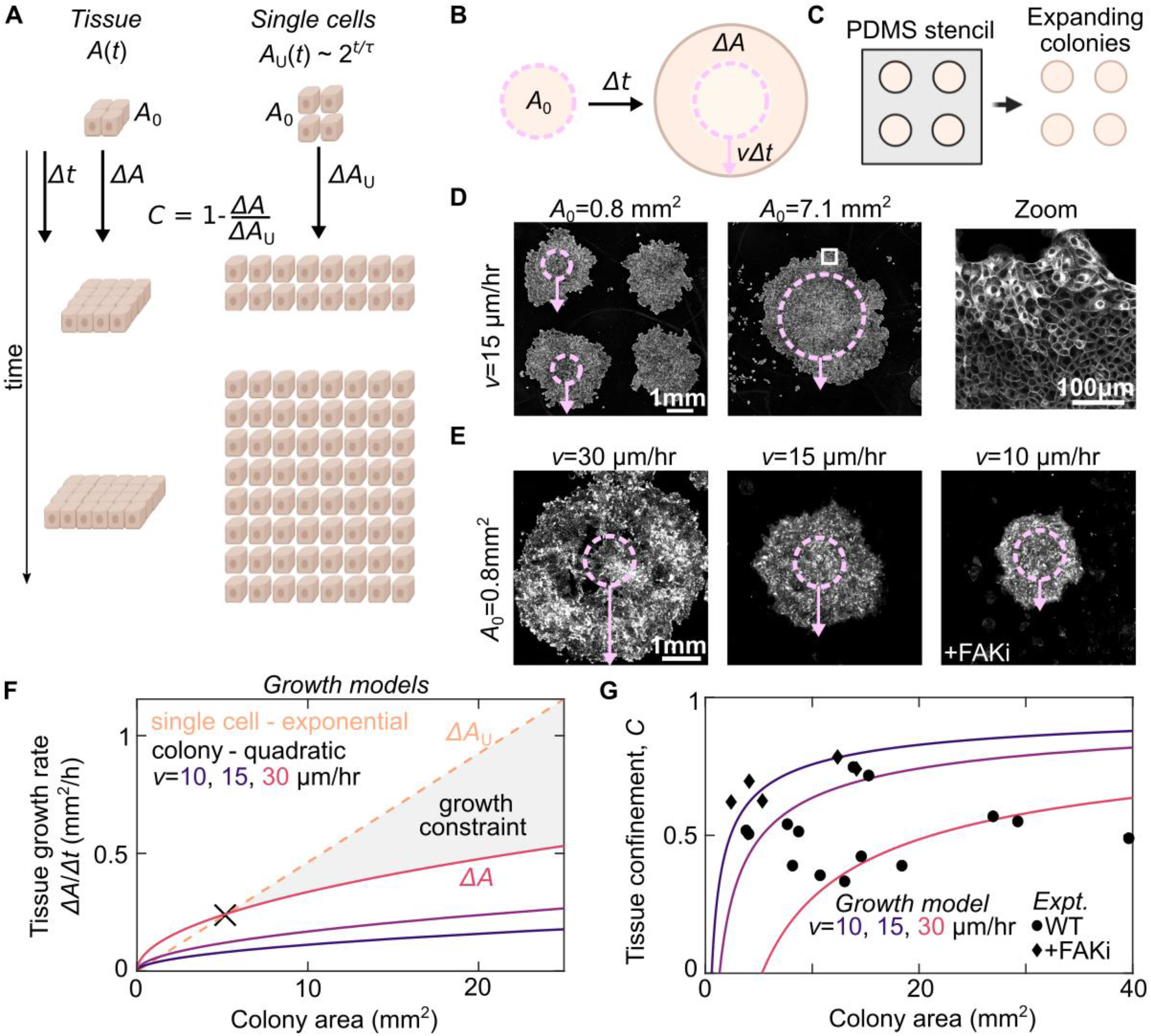
Tissue confinement quantifies how tissue-scale growth dynamics constrain cell growth. (A) We consider the growth dynamics of a portion of multicellular tissue with time dependent size A(t) with that of comprised of isolated cells of equal initial size, A_0_. The population of individual cells grows exponentially *A*_u_(t)∼2^t/τ^. The confinement, *C*, is measured by comparing the relative growth rates. (B) Schematic of expanding colony system. Cell colony of initial size *A*_0_ expands at a constant radial velocity *v* such that, for a given time interval *Δt*, the radius increases by *vΔt* (C) Cells are initially seeded into a PDMS well of defined size *A*_0_ then this well is removed to initiate colony expansion (D) Cell Mask Deep Red staining in expanding MDCK colonies with two different initial sizes at *Δt*= 48 hours after removing the barrier. Overlay shows the initial size *A*_0_ (dashed circle) and expansion *vΔt* (arrow). Zoom-in shows part of larger monolayer (white box). (E) CellTrace Far Red staining of expanding MDCK colonies with variation in expansion rate from variation in WT dynamics and under FAK inhibition (500nM PND-1186) Images are scaled differently for clarity and intensity is not comparable in these images. (F) Quadratic growth models of colonies as a function of tissue area and expansion rate compared with a model of the exponential rate for single cells growing at the experimentally measured proliferation rate 15 hours. (F) plot of the tissue confinement parameter, defined in the main text, calculated from comparing the model of colony growth rate to the model of single cell growth rate. Curves show tissue confinement as a function of tissue area and expansion rate. Black points show confinement measurements from the experimentally determined area growth rate of expanding colonies at *Δt* = 48 hours (see methods).

We explored this framework using model tissues comprised of large, circular colonies of MDCK cells, chosen for their well-characterized growth dynamics and the facility of controlling their size (Heinrich et al., 2020; Puliafito et al., 2012). Circular colonies of variable size A_0_ from ∼1-7 mm^2^ were formed by seeding the cells in a Polydimethylsiloxane (PDMS) stencil atop a glass coverslip (Fig. 3B&C). The stencil was then released to allow for colony expansion for Δt=48 hours (Fig. 3D). Importantly, the colony expansion is determined by radial migration speed *ν* of cells at the periphery to increase the colony radius by *ν*Δ*t* (Fig. 3B, arrow). Consistent with previous studies, the radial expansion rate was independent of colony size (Fig. 3D, Fig. S5). This results in a growth rate that scales quadratically with colony area (see Methods). For a given colony area, variation in *ν* provides additional control over colony growth rate (Fig. 3E). Under control conditions, *ν* varied between experiments from 15-30 μm/hr (Fig. S5). This tissue growth rate was not impacted when inhibiting cell division but was reduced by inhibiting cell migration by a focal adhesion kinase inhibitor (FAKi) (Fig. S5). Thus, variations in both the initial colony size and expansion velocity provide a wide range of tissue growth rates.

We generated model curves of the expected colony growth rates for the ranges of areas and edge velocities observed experimentally (Fig. 3F, solid lines). These rates were then compared to that expected for exponential growth of single cells, using the experimentally measured doubling time of 15 h (Fig. 3F, dashed line). For small areas, exponential growth is smaller than the growth rate of the expanding colony (Fig. 3F); here tissue-scale growth dynamics do not constrain cell proliferation. This is the behavior for SC. By contrast, for large areas, expanding colony growth rates become substantially lower than the exponential growth of single cells (e.g. Fig. 3F, gray shaded region). Here, tissue growth dynamics constrain cell proliferation. The colony area when exponential growth and quadratic growth rates are equal demarks the transition between these two regimes and, here, *C* = 0 (Fig. 3F, black X). For each given growth model, the confinement *C* is plotted as a function of colony area (Fig. 3G, lines). Confinement was determined from experimental data with varying *A*_0_ and *ν* (Fig. 3G, points). These experimental conditions then provide a means to systematically explore cell behavior in varied *C* from <0.25 to ∼0.8. Thus, our tissue confinement framework provides a quantitative method to assess how tissue-scale growth dynamics are expected to constrain growth of single cells.

To test the utility of this framework, we explored cell growth and signaling in tissues with varied levels of confinement. To query cell growth, we labeled the cells at t=0 with CellTrace, a fluorescent dye which reports on biomass production by its dilution (i.e. more growth leads to lower cell intensity) (see Methods). The total growth is determined from the CellTrace images by the ratio of intensity 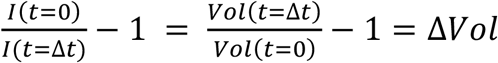. Since the growth rate of confluent monolayers is close to zero (Fig. 2B), the intensity of the confluent monolayer *I*_*C*_∼*I*(*t* = 0), can be used as the standard to compare to the intensity *I* of expanding colonies (Fig. 4A, *C*=1). This allows for determination of the growth rate by *I*_*C*_/*I* − 1. For Δ*t* = 48 hours, subconflucent cells show significant 10-fold CellTrace dilution consistent with the cells doubling every 15h (Fig. 4A, *C*=0). We then used expanding colonies with varied *A*_0_ to explore intermediate levels of confinement from 0.2 to 0.8. In smaller colonies with C=0.2, cell growth is already suppressed to <50% that of sub-confluent conditions. By C=0.6, growth is restricted to <10% that of the subconfluent cells and is suppressed to nearly zero by C=0.8 (Fig. 4A). Changes in growth can also be modulated by changing the edge velocity. In our experiments, we observed that differences in migration rate significantly impact the cell growth, as predicted by our modeling (Fig. 4C). In all conditions, the intensity is remarkably uniform across the tissue suggesting that the growth regulation mechanism is a tissue-scale phenomenon.

**Figure 4:**
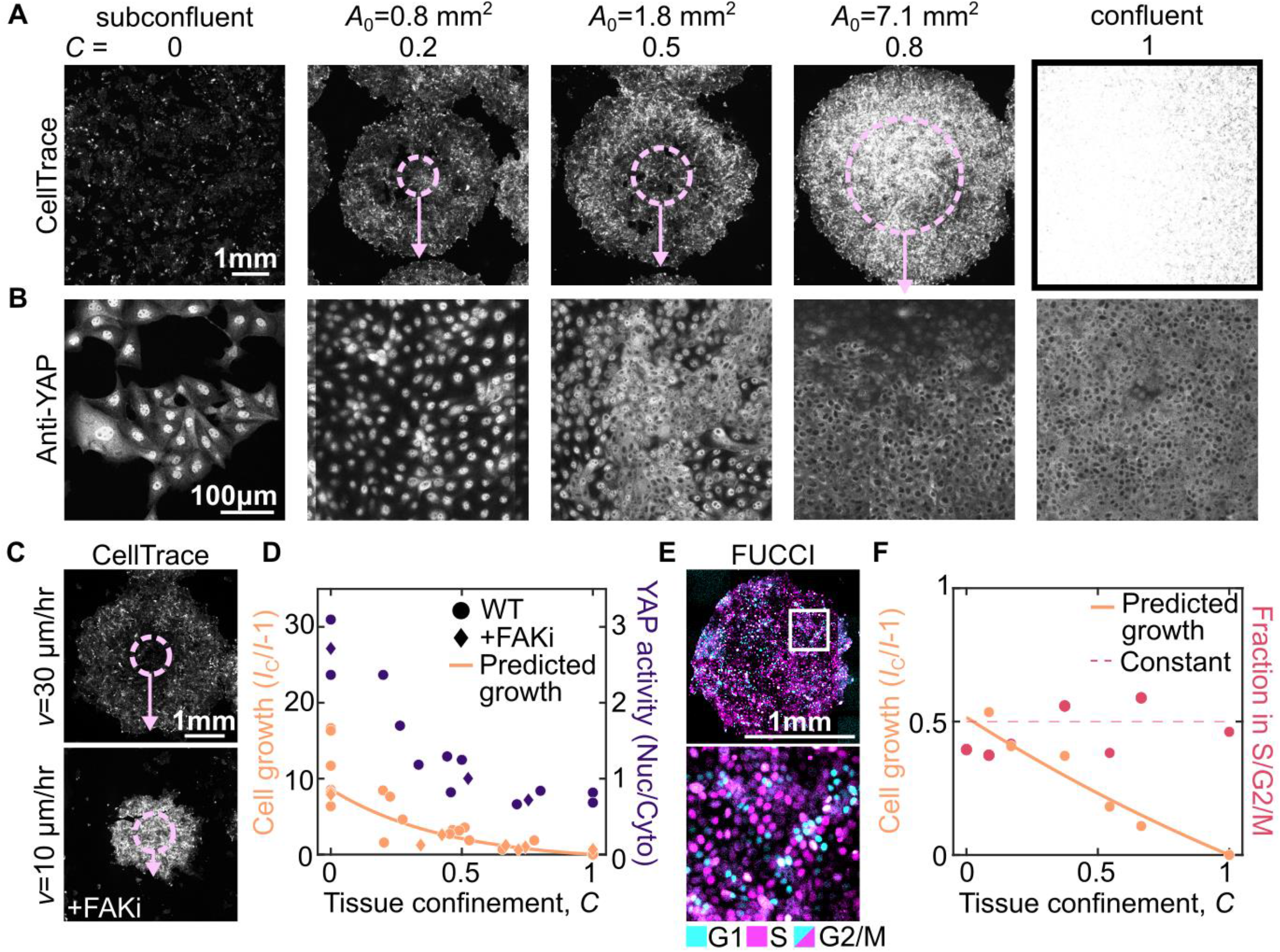
Tissue confinement determines cell growth rate and signaling. (A-B) Images of Cell Trace (A) and YAP (B) at *Δt* = 48h in expanding colonies with initial size, *A*_0_, varying from 0.8-7.1 mm^2^, subconfluent colonies and confluent conditions. In (A), overlay shows initial colony size A_0_ (dashed circle) and expansion *vΔt* (arrow). Note different scale bars in (A) and (B). (C) CellTrace images of expanding monolayers with *A*_0_=0.8mm at *Δt* = 48h in the presence and absence of 500nM PND1186 (+FAKi) to illustrate the effect of changes in *vΔt*. (D) Quantification of cell growth and YAP signaling at varying tissue confinements with *Δt* = 48h across multiple experiments. Each data point is an average of multiple colonies from one experiment (see Methods). YAP activity is determined by the nuclear to cytoplasmic ratio of anti-YAP intensity, quantified for >150 cells in each experiment. Growth is plotted against the time averaged confinement and YAP activity is plotted against the final confinement. Data are from 6 independent experiments (Cell growth) and 3 independent experiments (YAP activity) (D) MDCK FUCCI cells in expanding colony with *A*_0_=1.8mm^2^ and *Δt* = 12h. (E) quantification of cell growth and fraction of cells in the S/G2/M cell cycle states as a function of the average tissue confinement. Each data point is quantified from >3 colonies in a single experiment with A_0_=0.8mm^2^,1.8mm^2^,or 7.1mm^2^ and *Δt* = 12h. Data are from 2 independent experiments.

To query cell growth signaling under confinement, we also performed immunostaining against YAP. YAP is a transcription factor implicated in regulating cell proliferation during contact inhibition (Aragona et al., 2013; McClatchey and Yap, 2012; Zheng and Pan, 2019). When YAP is active, it is localized to the nucleus; when inactive, YAP is localized to the cytoplasm. We see in the conditions with lower confinement, a greater faction of YAP is localized to the nucleus, whereas around C=0.5 it becomes more localized to the cytoplasm (Fig. 4B). This suggests that YAP activity is regulated in response to changes in tissue confinement (Fig. 4B). Taking together, all our experimental data from colonies with varying size and edge velocity (Fig. 3G) reveal a systematic decrease in cell growth and YAP signaling as a function of confinement (Fig. 4D, points). Moreover, the predicted growth from the definition of tissue confinement is consistent with the experimental data (Fig. 4D, line). All of these data demonstrate the rapid suppression of cell growth at low levels of confinement.

At the onset of confluence, the cell division rate remains similar to subconfluent cells despite the increased confinement (Fig. 2B); this suggests that tissue confinement may not immediately affect the cell cycle. To query the cell cycle, we performed the expanding colony experiments with MDCK cells expressing the pip-degron fluorescent ubiquitination cell cycle indicator (FUCCI MDCK) (Grant et al., 2018) to measure the fraction of cells in S/G2/M in model tissues with varying levels of confinement. We restrict our analysis to short expansion times (Δ*t* = 12 h) before cells have reached the ME state and arrested the cell cycle. In contrast to cell growth (Fig. 4F, peach data), there cell cycle is insensitive to tissue confinement (Fig. 4F, red). Instead, the fraction of cells in S/G2/M is constant (Fig. 4F, dashed line). This data reveals the qualitatively different impact of confinement on cell cycle and growth. Thus, the transient uncoupling between cell cycle and growth observed in Fig. 2B is consistent with a rapid increase in confinement at the onset of confluence.

### A G1 sizer arrests the cell cycle in confined epithelium

We next explored how the cell cycle arrests in monolayers at the later stages of contact inhibition. After confinement reduces the growth rate, cell size decreases through successive cell division until the cell cycle becomes arrested (Fig. 2J). Previous work has shown that cell cycle regulation in isolated mammalian cells is cell size independent (Cadart et al., 2018) whereas it can be size-dependent for in *in vivo* epithelium (Xie and Skotheim, 2020). To examine if cell division is regulated by cell size in our data, we measured how the cell division rate varied as a function of cell size by tracking individual division events. We estimate volumes from the cell area multiplied by the typical cell height of 6.5±1.5 μm (Fig. S2, mean±S.D.). Above a volume of 1200 μm^3^, the cell division rate is independent of size (Fig. 5A). However, the division rate sharply decreases for smaller volumes. This trend is observed across a range of experimental conditions and two epithelial cell types, indicating it is a robust feature of cell size regulation in confluent epithelial tissue (Fig. S6). We then used FUCCI MDCK cells to look more closely at the cell cycle regulation for large and small cells. From the FUCCI data, we obtained the S/G2/M duration by tracking single cell trajectories and saw that the duration of the S/G2/M phases is ≈10 hours and independent of cell size (Fig. 5B). In the same data, the duration of the entire cell cycle was estimated by measuring the fraction of cells in each cell cycle phase (see Methods). We see that the cell cycle duration rapidly increases for smaller cells (Fig. 5B, purple data). The division rate in Fig. 5A can also be used to estimate cell cycle duration and shows a similar trend (Fig. 5B, dashed line). Together, these data indicate an increased duration of the G1 phase for smaller cells. This is consistent with previous results that size regulation occurs at the G1-S transition (Cadart et al., 2018; Murray and Hunt, 1993; Xie and Skotheim, 2020).

**Figure 5:**
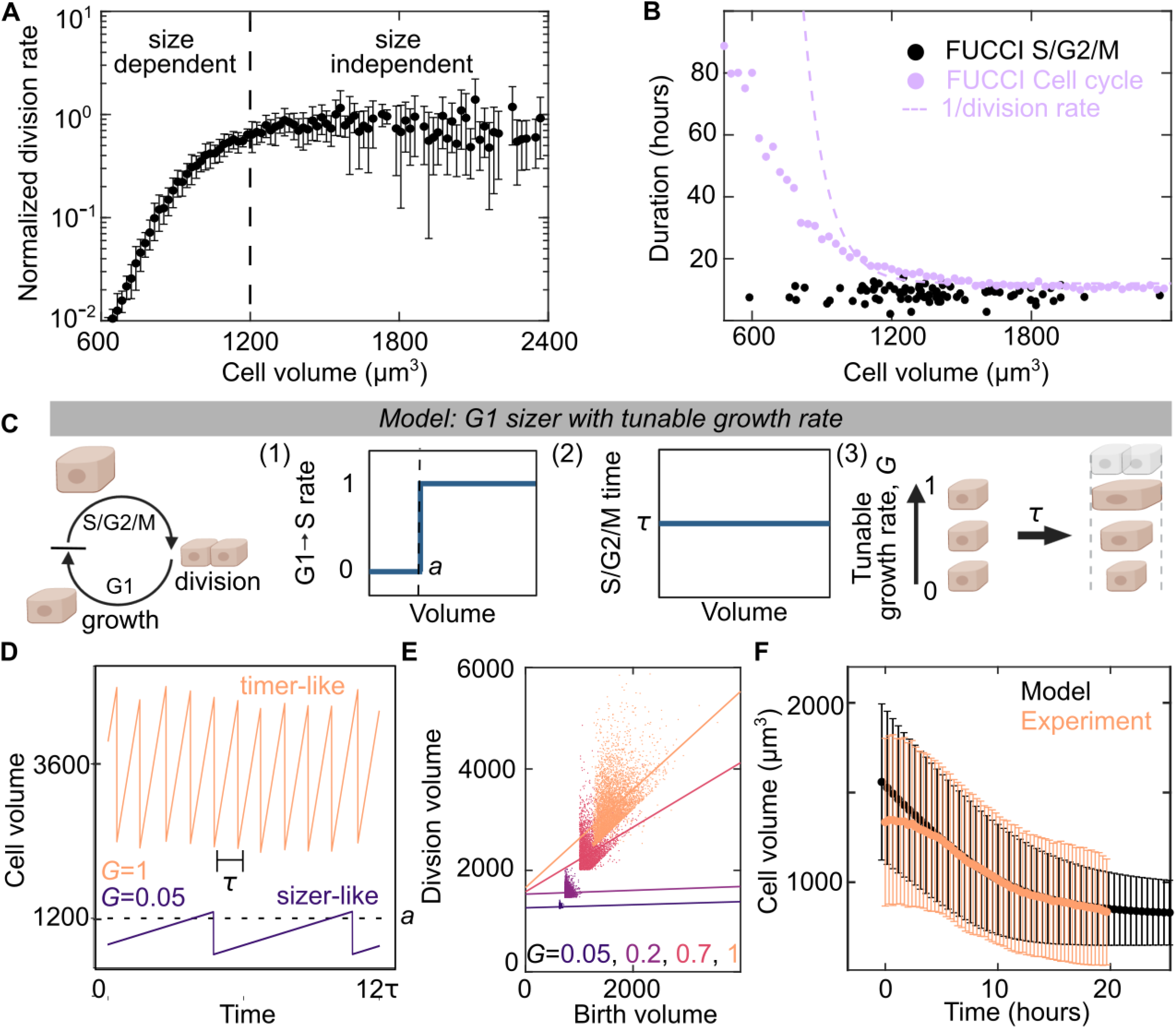
A G1 sizer arrests the cell cycle. (A) Normalized cell division rate as a function of cell volume in MDCK monolayers. Volume is calculated from cell areas multiplied by the average experimental height (see Methods). Data are averaged from 4 experimental replicates with >500 division events, error bar is standard deviation of experimental replicates (B) Duration of cell cycle (violet) and S/G2/M (black) for MDCK cells as a function of cell size. Dotted violet line is a fit to the data in A to extract the cell cycle duration. S/G2/M time are from N=82 trajectories for cell cycle. Cell cycle time data (violet points) are population averaged FUCCI measurements from 50 fields of view containing >100 cells from 1 experiment (see Methods) (C) Schematic of G1 sizer model – cells are simulated to grow at a constant rate, transition between cell cycle states and divide. (1) Cells transition rapidly from G1 to S only when above a critical volume *a*, (2) cells have a set S/G2/M duration equal to time τ that is independent of size, (3) cells have a variable growth rate, G, independent of the cell cycle (D) G1 Sizer models results of cell volume as function of time for two growth rates (G = 1 and G = 0.05) (E) G1 Sizer model results of cell volume at division as a function of birth volume for G=0.05, 2, 0.7 and 1. Dotted lines show a linear fit to the data (slope = 0,0,0.4,1). Each condition contains 400 simulation trajectories. (F) Cell volume as a function of time for experimental data from Fig. 2D (peach) and for G1 sizer model results (black). The onset of confluence occurs at t=0 for experiments. For simulations, this is models with a growth rate quench from 1 to 0 at t=-10h. Data is mean cell volume, error bars are the standard deviation. Each time point in the experiment is the average of >10000 cells from 30 fields of view. Each simulation time point is an average of >15000 cells from 35 simulations.

Motivated by this data, we developed a simple “G1 sizer” model of size-dependent exit from G1 (Fig. 5C)(Heldt et al., 2018; Xie and Skotheim, 2020). In the model, we simulate an ensemble of cells that grow at a constant rate, have two cell cycle phases G1 and S/G2/M, and divide into two daughter cells with half the mother volume. We added additional features based on experimental observations: (1) There is a sharp volume threshold of the G1-S transition rate and below this minimal size *a*, the transition rate is zero, (2) cells have an S/G2/M duration of *τ* = 10 hours independent of cell volume, and (3) a variable cell growth rate *G* that is normalized to vary from 0 for no growth to 1 for growth in the unconfined condition. Due to rule (2) the minimum cell cycle time is *τ* hours and cells will grow by a minimum of *Gτ* before each division. When *Gτ* ≫ *a*, cells are large compared to *a* and the G1-S transition proceeds quickly. In this regime, the cell cycle regulation is size independent (timer-like) with a time *τ* between cell divisions (Fig. 5D, *G* =1, Fig. S7). However, when the growth rate is suppressed such that *Gτ* << *a*, additional time is required to relieve the size constraint of the G1-S transition (1) (Fig. 5D, *G* =0.05, Fig. S7). In this regime, the cell cycle regulation is highly size-dependent (sizer-like).

Plotting the cell size at division as a function of cell size at birth for a range of growth rates shows that the model transitions smoothly from size-dependent to size-independent behavior as a function of growth rate (Fig. 5E, Fig. S8). Size-dependent cells divide at the same size and show no correlation between birth size and division size (Fig. 5E, dashed lines for *G*=0.05, 0.2). This contrasts with size independent cells which show a correlation between birth size and division size (Fig. 5E, dashed lines for *G*=0.7 & 1) (Amir, 2014; Cadart et al., 2018; Xie and Skotheim, 2020; Zatulovskiy and Skotheim, 2020). These different regulation mechanisms also occur in distinct cell size ranges consistent with previous work showing that large, rapidly growing, single cells are size independent (Cadart et al., 2018) and small, slowly growing cells, *in vivo* are size-dependent (Xie and Skotheim, 2020).

Having developed an understanding of the model at constant or near-constant growth rates, we tested if the model could also predict the cell size distributions found in the experiments of monolayer formation and maturation in Fig. 2B. We simulate monolayer formation in our model by a rapid quench of cell growth rate from 1 to 0 at t=0 and measure the cell size distribution over time (Fig. 5F). The simulation results (Fig. 5F, black) are consistent with those of the experiment (Fig. 5F, peach). This suggests that the G1 sizer model together with an understanding of how confinement impacts cell growth (Fig. 4F) is sufficient to explain transitions in size of isolated cells to those in epithelial tissue.

### Size-dependent Cyclin D degradation leads to cell cycle arrest

To investigate molecular mechanisms of size control, we took advantage of our Tet-inducible cell lines to manipulate cell size in confluent monolayers. We prepared monolayers at the onset of confluence with Tet-On p27 and Tet-On p27ck cells. At t=0 we added doxycycline (+dox) to induce expression. After 5 days, both monolayers are in a cell cycle and growth arrested ME state, but the +dox p27ck cells are 40% the size of +dox p27 cells (Fig. 2I; Fig. 6A). RNA sequencing revealed almost no differences in the steady-state transcriptome of these samples (Fig. 6B). However, close examination revealed several weak signatures (Fig. S9) including a slight downregulation of cyclin D1, 2, 3 in the smaller contact inhibited cells (Fig. 6B, inset). The G1/S transition has a steep dependence on cyclin D concentration (Fan and Meyer, 2021) so it is possible that small changes in cyclin D concentration are sufficient to arrest the cell cycle. Furthermore, cyclin D is strongly post transcriptionally regulated by degradation (Alao, 2007). To check if this difference in RNA abundance leads to changes in protein levels, we looked at cyclin D1 (cyD1) protein levels via immunofluorescence in ME formed with Tet-On p27 cells (+dox p27) and co-cultures including Tet-On p27ck cells (+dox p27/p27ck). Since p27 is known to interact with cyclin D, we tested that its overexpression did not change the cyclin D1 levels by inducing p27 after ME formation (Delay +dox p27). Interestingly, we found significant difference in cyclin D1 abundance, with nearly undetectable levels in cells with a nuclear area <100 μm^2^ (Fig. 6 C; Fig. S10). We observed that the intensity of CyD1 drops rapidly with decreasing cell size, measured by the nuclear area (Fig. 6D). The same trend was seen for all monolayer preparations, suggesting that the cyclin D1 level is regulated by a size-dependent pathway. This suggests that in addition to transcriptional changes in cyclin D, additional size-dependent post transcriptional regulation occurs, possibly as reported in other contexts (Alao, 2007; Masamha and Benbrook, 2009).

**Figure 6:**
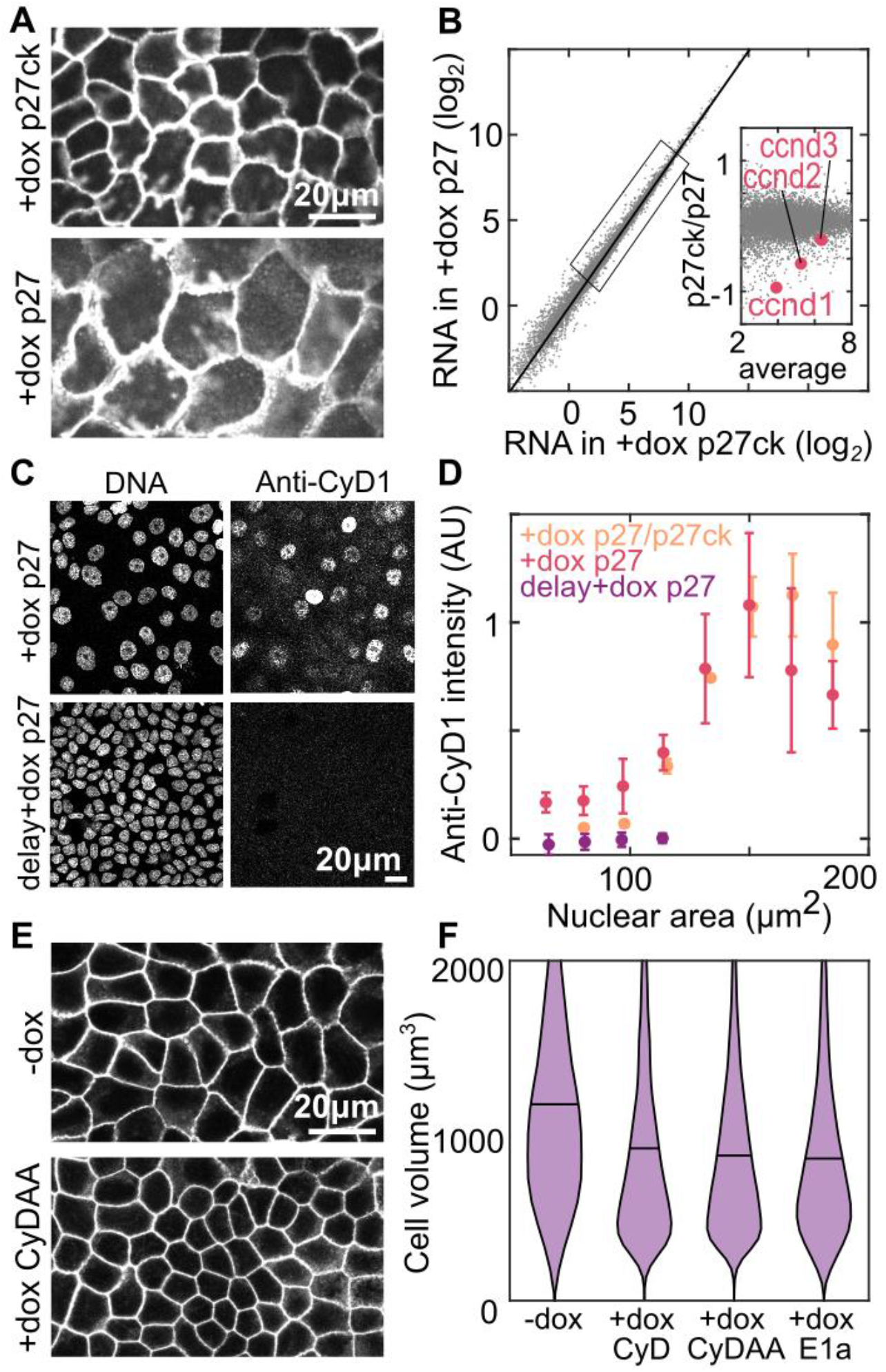
Low cyclin D causes cell cycle arrest in small cells. (A) Cell membranes of Tet-On p27 and Tet-On p27ck MDCK cells in ME+4 days; dox added at t=0h. (B) RNA sequencing data from monolayers prepared in A. Data are averaged transcripts per million from 3 experimental replicates. Inset: zoom in of genes in indicated box, cyclin D genes expression levels are highlighted in red. (C) DNA staining and anti-Cyclin D1(CyD1) immunofluorescence staining in MDCK monolayers at ME+3 days. Dox is added at either t=0 (+dox p27) or at t=ME+2 days (delay +dox p27). (D) Quantification of anti-CyD1 intensity from immunostaining data as a function of nuclear area. Intensity is normalized in each experiment to a maximum value of 1. P27/p27ck are monolayers with a mixture of Tet-On p27 and Tet-On p27ck cells. (N_+dox_=4564(3) N_Delay+dox_=7515(2) N_p27/p27ck_= 11080(4)) (E) Labeled cell membranes in Tet-On Cyclin D1 T286A T288A (CyDAA) MDCK monolayers at ME+3 days without dox (-dox) or with dox added at t=0 (+dox CyDAA). (F) Cell volume measured in resuspended Tet-On Cyclin D1, Tet-On Cyclin D T286A T288A, and Tet-On 12sE1a cells at ME+3 days without dox (-dox) or with dox added at t=0 (+dox CyD, +dox CyDAA, +dox E1a). (N_-dox_=13513(5) N_CyD_=7240(2) N_CydAA_= 9092(3) N_E1a_= 6457(4))

To test if decreased cyclin D levels are required to arrest the cell cycle, we overexpressed cyclin D1 in contact inhibited cells. We used Tet-On Cyclin D1-GFP (CyD) or Tet-On Cyclin D1 T286A T288A-GFP (CyDAA, a degradation resistant mutant) cells and induced the expression of additional cyclin D1 at OC. We then looked at the cell size 3 days later, after it had reached a plateau. We observe that overexpression of either CyD or CyDAA leads to decrease in minimal cell size in ME, compared to control (-dox) (Fig. 6E,F). Therefore, restoring Cyclin D1 in small cells is sufficient to initiate the cell cycle. This suggests that the depletion of cyclin D is necessary for size-dependent arrest of the cell cycle. We also overexpressed the viral oncoprotein E1a which is known to bind and inactivate Rb pocket proteins and activate the G1/S transition (Whyte et al., 1989). Cells that overexpressed E1a also showed decreased size, suggesting that cyclin D depletion arrests the cell cycle by inhibiting the G1/S transition (Fig. 6F).

### Cell cycle arrest occurs near cell size minimum set by the genome size

We next wanted to understand why the cell cycle normally arrests at a volume of ∼1000 μm^3^. A possible constraint on cell size comes from the volume occupied by the genome. As the cell size decreases, the nucleus gets smaller and chromatin gets more compact (Viana et al., 2021). A simple estimate suggests low chromatin concentrations ∼5% by volume in an average subconfluent mammalian cell (Vol_nuc_∼1/3Vol_cell_ ∼800um^3^ vs Vol_genome_ ∼ 40um^3^) (Milo and Phillips, 2015). However, the concentration would increase several fold as cell size reduces and the total chromatin per cell remains constant. Previous measurements of chromosome size by TEM and AFM have shown that the volume of a full set of chromosomes are approximately 50-100 μm^3^ (Fritzsche and Henderson, 1996; Heslop-Harrison et al., 1989) and 50% chromatin by volume (Ou et al., 2017), thus, setting a lower limit on cell size. When we stained both the DNA and cell membrane, we observed that the abnormally small Tet-On CyDAA cells appear to have an unusually large nucleus relative to the cell size (Fig. 7A). This was surprising given that there is typically a tight scaling relationship between cell size and nuclear size (Viana et al., 2021). Comparing the cell volume against the nuclear volume for epithelium prepared under our previously described conditions, we observe this scaling relationship except in the Tet-On CyDAA cells in the presence of doxycycline (Fig. 7B, peach). Instead, we observe that the ratio of nuclear to cell size is rapidly increasing as cell size decreases below 1000 μm^3^ (Fig. 7C) and approaches the regions where DNA compaction exceeds the chromosomal values or where nuclear size would exceed cell size (Fig. 7B, gray, Fig. 7C gray). We hypothesized that increasing chromatin concentration could disrupt normal chromatin function, leading to DNA damage. In abnormally small cells we observed phospho-H2A.X foci indicating locations of DNA damage (Fig. 7D, E). This suggests that normal cells arrest near a cell size minimum but outside the range where DNA damage occurs frequently. DNA damage is known to arrest the cell cycle through Rb/Cyclin D independent mechanisms (Shaltiel et al., 2015) preventing further size reduction. Therefore, in epithelia, proliferative homeostasis is maintained by an interplay between cell growth in proportion to tissue constraints and cell size-dependent G1/S regulation which arrests cell size near a minimum (Fig. 7F).

**Figure 7:**
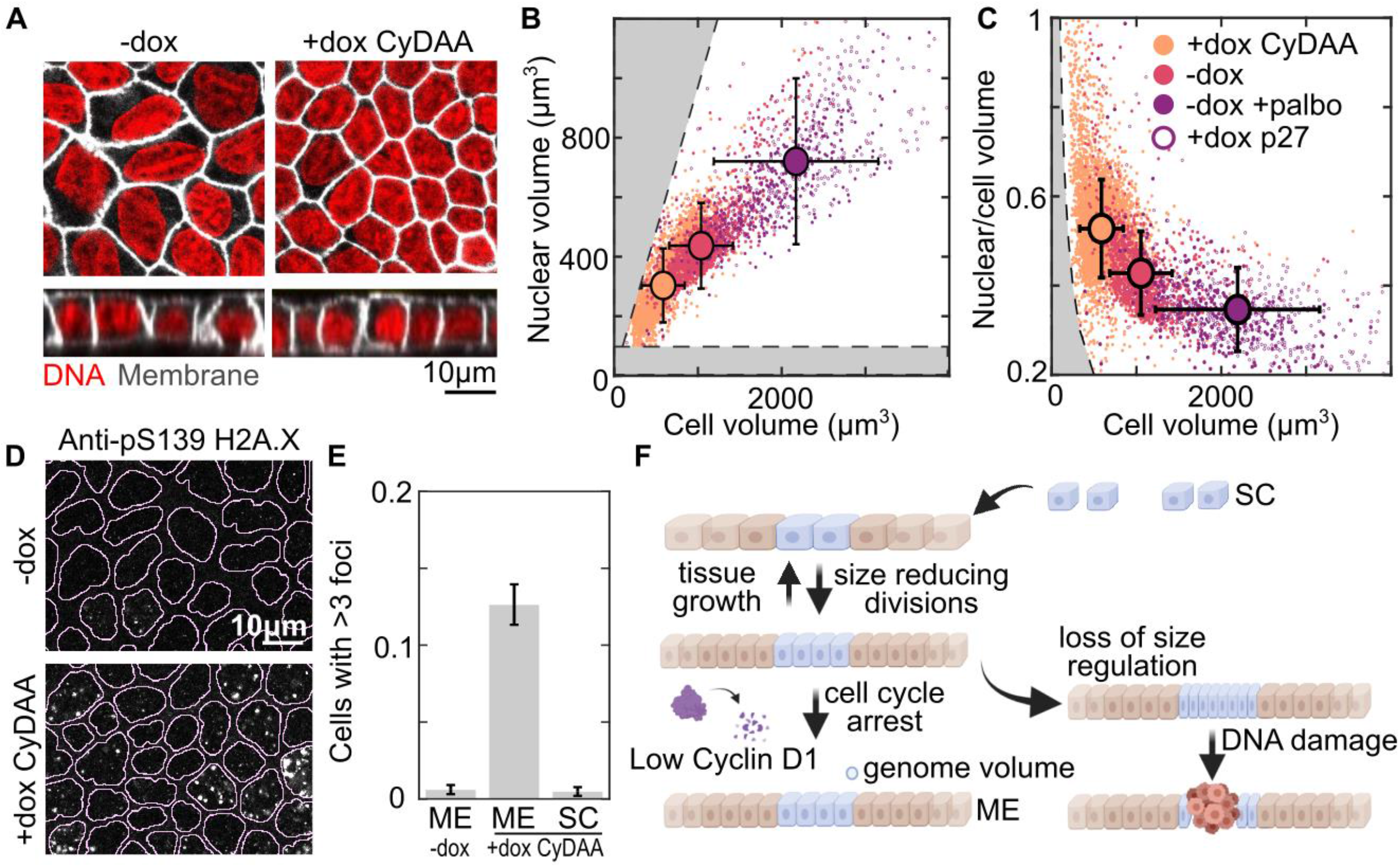
Cells arrest near a minimum size set by genome volume. (a) Nuclear (red) and membrane (white) staining of Tet-On Cyclin D1 T286A T288A MDCK cell monolayers at ME+3 days without dox (-dox) or with dox added at t=0 (+dox CyDAA)(B-C) Plots comparing nuclear volume and cell volume measured from 3D imaging in conditions from A (-dox, +dox CyDAA) or in cell cycle inhibited conditions (-dox +palbo) or Tet-On p27 cells (+dox p27). +palbo is 1µM Palbociclib. B shows the correlation between the cell and nuclear volumes while C show the ratio of these volumes. Gray regions indicate when nuclear volume is larger than cell volume (B-top left) or when chromatin density is larger than chromosomes (B-bottom, C-left). Error bars show standard deviation of data(N_-dox_=1405(2) N_+dox CyDAA_=4207(2) N_-dox +Palbo_= 390(1) N_+dox p27_= 324(1)) (D) Monolayers in the same conditions as A immunostained for pS139 H2A.X (yH2A.X). Lines overlayed on images show nuclear segmentation from DNA staining. (E) Quantification of pS139 H2A.X foci from conditions in D. Bar is the mean fraction of cells with >3 foci between 3 experimental replicates. Error bar is standard deviation of different experiments (N_ME-dox_=5813(3) N_ME+dox CydAA_=9127(3) N_SC+dox CydAA_= 2470(3)) (F) schematic summarizing cell size regulation in epithelium.

## DISCUSSION

While growth and division are coupled in single cells, we observe their regulation is uncoupled in epithelial tissue. The tissue growth dynamics regulate cell growth whereas cell division is regulated solely by cell size. Differences in these two rates drive changes in cell size depending on the tissue environment. In single cells, the environment places no constraint on cell growth leading to high growth rates and large cell size. In contrast, in mature epithelia, tissue growth rates are low and reduce cell size to a minimum. In this regime, cell size regulation is critical for maintaining cell homeostasis and preventing DNA damage. The consistency in cell size distributions across diverse epithelial cell types (Fig. 1B) suggest that our models should be broadly applicable to understand contact inhibition and cell size regulation across diverse biological systems.

Canonically, growth factor signaling is thought to be the main pathway to control and coordinate proliferation (Liang et al., 2017; Schlessinger and Ullrich, 1992). We identify an independent role for tissue confinement in controlling cell growth. While we exploited model tissues with a particular type of growth dynamics driven by edge migration, these ideas can be easily extended to arbitrary systems so long as the tissue growth dynamics can be readily characterized (Fig. S11). Tissue growth is driven by diverse processes, including migration, tissue buckling or mechanical stretch and occurs in response to cell turnover (Cowin, 2004; Guillot and Lecuit, 2013), all of which would reduce tissue confinement. Previous work has implied that cell growth and YAP/TAZ signaling in epithelium are regulated by mechanical stress (Irvine and Shraiman, 2017; Pan et al., 2016; Puliafito et al., 2012; Shraiman, 2005; Streichan et al., 2014). Our framework provides a means to isolate the roles of physical constraints on cell growth regulation. Importantly, confinement is a geometric quantity readily determined from timelapse microscopy. Our data show that confinement is a strong predictor of YAP/TAZ activity, demonstrating the utility of our model to study the mechanisms underlying epithelial growth control. However, future work is needed to determine the relationships between tissue confinement, growth and mechano-transduction.

Our observation of a transition between size-dependent and independent behaviors in epithelia may explain prior observations of size regulation in mammalian cells (Cadart et al., 2018; Xie et al., 2022; Xie and Skotheim, 2020). In our computational model we find that a size-dependent G1/S transition gives rise to both sizer-like and timer-like behaviors of cell size regulation at low and high growth rates, respectively (Fig. 5E). In future work, such a cell cycle model may be extended to other cell types or non-mammalian systems which show uncoupling between growth and division like Chlamydomonas or cyanobacteria (Li et al., 2016; Liao and Rust, 2021). Furthermore, the molecular mechanisms of G1 sizer regulation have remained elusive (Xie et al., 2022; Xie and Skotheim, 2020; Zhurinsky et al., 2010). By experimentally manipulating cell size, we showed that cyclin D1 regulation underlies G1 sizer behavior in epithelium. Cyclin D1 is strongly post transcriptionally regulated by degradation (Alao, 2007), suggesting that upstream kinase localization or activity may function in the size sensing pathway. Our observations may lead to future work to connect cyclin D1 regulation directly to cell size sensing.

Finally, below the minimal size set by cyclin D1 regulation, significant DNA damage occurs suggesting an important role of size regulation in maintaining cell homeostasis. Cancers which are driven by mutations in genes implicated in cell size regulation, such as small cell cancer, may show similar DNA damage leading to additional mutations. Alongside recent work which shows that very large cells become nonfunctional (Cheng et al., 2021; Lanz et al., 2021; Neurohr et al., 2019), the lower bound set by the genome size establishes a range of cell sizes for viable diploid mammalian cells from ∼200-10000 µm^3^, similar to the range observed across different cell types (Milo and Phillips, 2015). Overall, our understanding of the proliferative behaviors in epithelium provides a new basis for studying development, homeostasis, and disease in complex epithelial tissues across diverse biological contexts.

## Acknowledgments

Graphical abstract, Fig. 2C, 3A-C, 5C, 7F created with BioRender.com. This work was funded by NIH RO1 GM104032

## Author contributions

Conceptualization, J.D., M.L.G.; experimental methodology, J.D., M.L.G and L.J.H; modeling methodology, M.J.F., A.M.; formal analysis, J.D. and M.J.F.; investigation, J.D. and M.J.F.; resources, J.D. and M.J.F; writing – original draft, J.D. and M.L.G.; writing – review & editing, J.D., M.L.G., M.J.F., A.M, and L.J.H.; visualization, J.D. and M.J.F.; supervision, M.L.G., A.M. and L.J.H.; funding acquisition, M.L.G. and A.M.

## Declaration of interests

The authors declare no competing interests.

## Data Availability

RNA sequencing data will be deposited at the Gene Expression Omnibus prior to publication. Plasmids will be deposited to addgene prior to publication. Analysis and simulation code will be published to Github prior to publication. Image data and histology analysis will be deposited to figshare prior to publication.

## Lead Contact Statement

Further information and requests for resources and reagents should be directed to and will be fulfilled by the lead contact, Margaret Gardel.

## MATERIALS & METHODS

## REAGENTS

PND1184 and (3-Aminopropyl)trimethoxysilane were purchased from Sigma-Aldrich (Saint Louis, MO). Cell trace purchased from Invitrogen (Waltham, MA). Glutaraldehyde purchased from Electron Microscopy Sciences (Hatfield, PA), BD Collagen I, rat tail was purchased from BD Biosciences (San Jose, CA). 1X PBS, 1X DMEM, Fetal Bovine Serum (DMEM), l-glutamine, Penicillin, Streptomycin, Trypsin EDTA were purchased from Corning Inc. (Tewksbury, MA), TBS, MnCl, NaOH were purchased from Fisher Scientific (Hampton, NH), Palbociclib was purchased from Cayman Chemical (Ann Arbor,MI) Anti-cyclin D1 ((E3P5S) XP® Rabbit mAb 55506), Phospho-Histone H2A.X (Ser139) (20E3) Rabbit mAb #9718. purchased from Cell Signaling Technologies (Danvers, MA). Anti Yap purchased from Santa Cruz Biotechnology (Dallas, TX). Infusion and Lenti-X(tm) Tet-On® 3G Inducible Expression System purchased from Takara Bio (San Jose, CA). Janelia Fluor 646 halotag ligand, Janelia Fluor 549 halotag ligand and Fugene HD purchased from Promega (Madison, WI)

## CELL CULTURE & LINES

All cells were maintained at 37C and 5% CO2. Cells were passaged using 0.25% trypsin EDTA every 2-3 days. Cells were checked for mycoplasma by Hoechst staining. MDCK, CACO-2 and MEF cells were cultured in dulbecos modified eagle medium (DMEM) high glucose supplemented with 2mM L-glutamine and 10% FBS. HaCaT cells were maintained in low calcium high glucose DMEM prepared from calcium-free DMEM powder (#09800; US Biological) supplemented with 40 μM calcium-chloride, 2 mM L-glutamine and 10% calcium depleted FBS using Chelex-100 (Sigma-Aldrich). RPE-1 cells were maintained in 1:1 high glucose DMEM:F12k supplemented with 2mM glutamine and 10% FBS. Tet inducible gene expression was done with 200ng/ul doxycycline in all indicated experiments (+dox).

HaCaT cells were provided by Yu-Ying He (University of Chicago). Caco-2 cells (HTB-37) and HEK293T cells (CRL-3216) were acquired from ATTC. MDCK cells were provided by James Nelson. RPE-1 cells were provided by Wallace Marshall. MEFs were provided by Mary Beckerle. Stargazin-halotag Caco-2 and MDCK cells were produced by lentiviral infection of CACO-2 and MDCK cells by a WPT-Stargazin-halotag construct packaged in 293T cells by a second generation lentiviral system with pHR1-8.2-deltaR and a VSV-G pseudotyping plasmid (gifts from M. Rosner). Viral supernatant was collected at 24, 48 and 72 hours after transfection then concentrated ∼30x using Amicon Ultra-15 Centrifugal Filter Unit (100kDa) or concentrated ∼30x by peg precipitation (Marino et al., 2003). Cells (∼50,000 cells a in 6cm diameter dish) were treated overnight with 300ul of concentrated virus in 2ml of media supplemented with 8µg/ml polybrene. Positive cells were isolated using a cell sorter. FUCCI MDCK cells were produced by lentiviral infection with virus of pLenti-PGK-Neo-PIP-FUCCI packaged and infected the same way. Cells were then selected using 800 µg/ml G418. pLenti-PGK-Neo-PIP-FUCCI was a gift from Jean Cook (Addgene plasmid # 118616 ; http://n2t.net/addgene:118616 ; RRID:Addgene_118616). Tet mEmerald-P27 1-176, snaptag-P27ck 1-176, Cyclin D1-mEmerald, Cyclin D1 T286A T288A-mEmerald, mKate-T2a-12sE1a cells were produced using the Lenti-X Tet-On 3G system (Takara Bio). DNA above were subcloned into the Tre3g vector. Lentiviral particles were packaged in 293T cells transfected with pHR1-8.2-deltaR and a VSV-G. Cells were infected with lentivirus with both the EF1a-Tet-on-3g and Tre3g plasmids above using the infection protocol above then selected using 2 µg/ml puromycin and 800 µg/ml G418. Plasmids used in this study will be available on addgene.

## SAMPLE PREPARATION & MEASUREMENT

### Epithelial monolayer cultures

For all experiments unless indicated otherwise, monolayers were formed on 2mg/ml collagen I gels (∼200um thick) formed on top of a coverglass substrate (see Collagen gel substrate preparation section for more detail). For monolayer samples to reach the OC state after ∼12 hours and ME at ∼36 hours cells were seeded onto collagen gels at high density (∼80,000 cells/cm2). For SC samples a low density of cells (∼8,000 cells/cm2) were plated on the same substrates and cells were measured before reaching OC. Cells were added on top of the gel in a volume of 100-200ul and allowed to adhere for 5-10 minutes before adding 1.5ml to the culture dish containing the coverslip and gel. Culture media was changed once each day.

### Expanding colony assay

Expanding colonies were prepared using published methods (Heinrich et al., 2020). 4.4 grams of 10:1 PDMS (silgard) was cast in a 10cm petri dish and cured at 70C overnight. A piece of PDMS ∼20×20 mm was cut out then a set of holes was cut into the PDMS using a leather hole punch of 1mm,1.5mm or 3mm (Nuhank 0795787181775). The PDMS was washed in 70% ethanol for 5 minutes repeated 3 times then milli-Q water 3 times and allowed to dry. Cover slips were coated with collagen 1 by incubating them on a drop of 0.2mg/ml collagen in 0.02M acetic acid for 1 hour. Coverslips were washed with 1xPBS 3 times then with MQ water 3 times and allowed to dry completely. Dry PMDS and coverslips were stuck together ensuring that no air bubbles remain between the surfaces. Cells were seeded in the well (2000 cells/mm2) and allowed to adhere for 5-10 minutes before adding 2ml of media to the petri dish. Colonies were left overnight then the PDMS was removed to allow colonies to expand. The initial colony size under these conditions was measured and used for subsequent analysis. Each experiment included a subconfluent and confluent sample as exponential and non-growing controls to ensure that results could be compared across experiments. After the desired time delay samples were either fixed and imaged or fixed, permeabilized, immunostained and imaged according to methods below.

### Cell Trace labeling

Cells were labeled using cell trace according to the manufacturer’s protocol. Cells were resuspended in PBS (10^6^ cells in 1 ml) and cell trace was added at 1μM for 15 minutes at 37C. Then cells were pelleted and resuspended in media and either directly used for experiments or cultured under normal conditions for 1 day before use.

### Immunostaining

For halotag labeling, 30nM halotag JF 646 or 549 solution was added for 1 hour before fixation. Just before fixation cells were washed once in 1xPBS. Cells were fixed in 4% PFA in 1xPBS for 15 minutes at room temperature. Cells were blocked and permeablized in 1xTBS, 0.3% triton-X 100, 2%BSA solution for 1 hour. Antibody solutions were prepared in 1xTBS 0.3% triton-X 100 2%BSA using 1:400 anti-Cyclin D1, 1:100 anti-YAP, 1:400 anti-Phospho-Histone H2A.X. Samples were incubated in primary antibody overnight at 4C. Samples were washed 3 times in 1xPBS then incubated in 1xTBS 0.3% triton-X 100 2%BSA and secondary antibody for 1 hour. In conditions with DNA staining 1x SPY650 DNA was added during the secondary staining step according to the manufacturer’s protocol at 1x concentration. Samples were washed 3 times for 5 minutes in 1xPBS then mounted on a slide in prolong gold antifade mounting media – non curing (Invitrogen), sealed and imaged.

### Fluorescence microscopy

For time lapse imaging cells were imaged on an inverted epi-fluorescence microscope (Nikon TI-E, Nikon, Tokyo, Japan) with a 20x plan flour multi-immersion objective. images were acquired at 10-minute intervals in GFP, 642 and transmitted light channels using standard filter sets (Ex 490/30, Em 525/30, Ex 640/30, DAPI/FITC/TRITC/cy5 cube) (Chroma Technology, Bellows Falls, VT). For halotag labeling, 30nM halotag JF 646 or 549 solution was added for 1 hour before imaging. Samples were mounted on the microscope in a humidified stage top incubator maintained at 37C and 5% CO_2_. Images were acquired on either a Photometrics Coolsnap HQv2 CCD camera (Photometrics, Tucson, AZ) or Andor Zyla 4.2 CMOS camera (Andor Technology, Belfast, UK).

Cell volume measurement samples in resuspension were imaged on an inverted spinning disk confocal microscope (Nikon TI-E) with laser lines at 491, 561 and 642 and suitable emission filters (Chroma Technology). Images were acquired using a 40x plan fluor oil immersion objective (NA 1.3) and Andor Zyla 4.2 CMOS camera (Andor Technology, Belfast, UK). Images were acquired at room temperature within 1 hour of cell resuspension.

All other imaging was done using a point scanning confocal microscope (Ziess Airyscan LS980) with laser lines at 491,561,642 and an adjustable emission filter suitable for fluorophores that were imaged. Cell trace images were acquired using a 5x air objective (NA 0.16), YAP images were acquired using a 20x air objective (NA 0.8), Immunostaining images were acquired using a 40x oil immersion objective (NA 1.3).

### RNA Sequencing

Cells were treated with 1um Palbociclib for 16 hours then replated to make 3 monolayers from each condition. All monolayers were cultured together in a 10cm petri dish with 10ml of media containing 100ng/ul doxycycline. Media was replaced each day for 5 days. Then 2 monolayers from each condition were lysed, pooled and total RNA was collected using a NucleoSpin RNA kit (#740955; Macherey-Nagel). 1 monolayer from each condition was labeled with JF 646 halotag and imaged, then cells were resuspended for volume measurements as described above. Volume distributions corresponding to each RNA seq experiment are available in Fig S10. RNA samples were submitted to the University of Chicago Genomics Facility. The sequencing facility performed QC measurements on RNA (RIN from 9.4-10), libraries were prepared using Oligo-dT mRNA directional primers, and sequenced using Illumina NovaSeq 6000 with ∼60M PE reads/sample. Alignments were made to the canine genome (Canis_lupus_familiaris.CanFam3.1) by psudeoalignment using Kallisto 0.46.1 (Bray et al., 2016). between 62.2 and 69.2% of reads were mapped with two technical replicates per experiment a total of N_p27_1_ = 28049196 N_p27_2_ = 28117815 N_p27_3_ = 28012202 N_p27ck_1_ = 28212157 N_p27ck_2_ = 27961694 N_p27ck_3_ = 27964407 reads. Data were then processed using iDEP 0.91(Ge et al., 2018) to measure differential gene experssion (“Wnt Target genes | The Wnt Homepage,” n.d.).

### AminoSilane Glutaraldehyde modification of glass coverslips

Glass coverslips were modified as previously described (Zhu et al., 2012). Coverslips were first cleaned by sonication in 70% and 100% ethanol solutions then dried with compressed air. We placed coverslips in a staining rack and submerged the rack in a solution of 2% (3-Aminopropyl)trimethoxysilane (APTMS) 93% propanol and 5% DI water for 10 minutes at room temperature while stirring. Staining racks were removed and washed in DI water 5 times then placed in a 37C incubator for 6-12 hours to allow the water to dry and amino-silane layer to cure. The staining racks were then submerged in 1% glutaraldehyde in DI water for 30 minutes while stirring. The coverglass was washed 3 times for 10 minutes in distilled water, air dried and stored at room temperature. Activated coverslips were stored under vacuum and used within 6 months of preparation.

### Collagen gel preparation

10x PBS, milli-Q water, a 5mg/ml collagen stock and 1N NaOH were mixed to generate a polymerization mix with 1xPBS and 2 mg/ml collagen at pH ∼7. 70uL of the polymerization mix was added on to a 22×22mm coverslip modified with aminosilane according to the protocol above and quickly spread to coat the surface using a pipette tip. Samples were transferred to a humidified incubator at 37C to polymerize for 20 minutes. After polymerization gels were washed 3 times in 1x PBS and it was verified that gels were still intact and adhered to the glass by a tissue culture microscope.

## CELL VOLUME MEASUREMENTS

### Resuspended

Cells were plated as monolayers for indicated times. Just before making volume measurements the cells were resuspended by adding 0.25% trypsin EDTA solution to the cells. Resuspending cells from ME conditions required partial physical disruption of the monolayer using a pipette tip. Cells were resuspended in 35ul of media containing 30nM JF646 and incubated for 5-15 minutes before adding to the sample prepared by sticking a coverslip to a glass slide using double stick tape. Cells were imaged immediately with spinning disk confocal microscopy. We verified that samples showed no changes in cell volume measured over time up to 1 hour and performed all measurements within this time window. Cells were imaged at the middle plane so that the radius of the cell could be measured. The cross-sectional area was used to estimate the radius which was used to calculate the volume of the cell. It was verified that this provided a comparable measurement to 3D segmentation of cells in the monolayer (Fig S1).

### 3D Images

Monolayers were stained using JF 549 halotag ligand and imaged using an airyscan LSM 980. Z-stacks spanning the height of the cell were imaged. The height of the monolayer was determined at each point by identifying maxima of the intensity corresponding to labeling at the top and bottom membrane of the cell. The membrane label averaged across the middle 5 planes of the cell was used to determine the area of the cell in the XY plane and this value is multiplied with the average cell height contained within this region to give the cell volume. A similar process was repeated with nuclei images to measure the nuclear volume.

### Cross-sectional Images

3D images were acquired above and displayed as projections in the YZ plane. Images were opened in imageJ and the width and height of individual cells was measured from these cross sectional images. The same method was applied to histology sections where cells oriented perpendicular to the tissue section were first identified then the width and height of these cells was measured. The volume was estimated by the width squared times the height of each individual cell. These measurements are compared with 3D images for MDCK cells (Fig. S1)

## IMAGE ANALYSIS

### Segmentation

Images of cell membranes were segmented using custom MATLAB code. The main algorithm performs initial segmentation using the Phase Stretch Transform algorithm developed by the Asghari and Jalali (H and JalaliBahram, 2015). Phase stretch images were thresholded and skeletonized to obtain cell outlines. Broken edges in the skeleton were repaired using a modified implementation of edgelink developed by Peter Kovesi (“Peter’s Functions for Computer Vision,” n.d.). Segmentation code can be made available upon request or from github/gardelLab.

### Cell tracking

Cell tracking was performed using established particle tracking methods (Crocker and Grier, 1996). Cell centers were determined by taking the centroid of each segmented cell area generated as described above in Segmentation. The particle trajectories were compiled from these position measurements using SimpleTracker, a MATLAB function developed by Jean-Yves Tinevez (“simpletracker,” n.d.).

### FUCCI Analysis

Cells were imaged in GFP and RFP channels similar to above methods using timelapse imaging. Images of FUCCI markers and cell boundaries were segmented using Phase Stretch Transform in Matlab as described above. Each cell was identified using the cell boundaries and was determined to be GFP or RFP positive by measuring the intensity contained within the segmented images of each nuclear marker. The percent of cells in G1 was determined by taking the ratio of cells identified as only GFP positive to the cells identified as GFP positive, RFP positive and positive for both markers. To determine duration of S/G2/M phase the cell cycle state was measured along the cell trajectory and points where the cell switched from G1 to S and then back to G1 were identified. Then the time between these events was measured to give the duration. To determine the full cell cycle duration the fraction of cells in S and G2/M phase was measured and along with the S/G2/M phase duration was used to estimate the cell cycle duration by CC duration = S phase duration/S phase fraction (i.e. if 10% of cells are in S/G2/M which lasts 10 hours, cells spend 9 times longer on average in G1 and the duration of the cell cycle is likely 100 hours)

### Division rate measurement

We identified cell divisions by finding pairs of cells which appear adjacent to each other in a frame after both cells were not present in the previous frame. We further filter out cells which are not of similar size to one another. We confirmed by inspection that this gives us a subset set of cells which have divided in the previous frame with few false positives. We then find the mother cell by looking several frames back for a cell near the centroid of the pair of daughter cells. We compare the number of cells of a given size which are detected to divide compared to the total number of cells of that size to get a probability of division. This process is repeated for the entire time series with a total number of division events typically >500. The division rate is determined by the change in cell density over time to compute the overall rates

### Quantification of H2ax staining

Foci were segmented using a phase stretch transform-based method and an intensity threshold. The cell nuclei were segmented using methods above and the number of foci in each nucleus was quantified.

## MODELING

### Growth Models

Growth curves were generated from growth models of exponentially proliferating cells with doubling time τ (A(t) = 2^t/τ^, dA(t)/dt = log(2)/τ * 2^t/τ^ = A(t)*log(2)/τ) and of a circle with an expanding radius (r(t) = v*t, A(t) = π*v^2^*t^2^, dA(t)/dt = 2*π*v^2^*t = 2*v*sqrt(π*A)). Confinement curves are calculated from the ratio of these rates as defined in the main text.

### G1-Sizer Model

Our phenomenological model of cell size control is a “G1 Sizer”, which posits that exit from G1 is controlled by a size-dependent function. Based on the sharp drop-off in the G1 exit rate seen in the log-plot of experimental data in Fig. 5A, we assume that the rate is a constant *k* ≫ 1 above a critical size *a* and 0 below that size. Following G1-exit, division proceeds in time *τ*.

As discussed in the main text, this model has two regimes: one of slow growth (*Gτ* ≪ *a*), and one of fast growth (*Gτ* ≫ *a*). Switching to non-dimensional units where *a* = 1 and *τ* = 1, we can derive results for time-averaged single-cell quantities, including average area, average time between divisions, and confinement. This can be done for both the fast- and slow-growth limits. In the following expressions, < > indicates an average over time for a single cell.

In the fast-growth limit with growth rate *G*, cells never interact with the size-threshold a, and hence have a constant division time set by the mean length of G1 plus the length of S/G2/M. This means a confinement of 0, and an average size *s* proportional to *G*. To summarize:

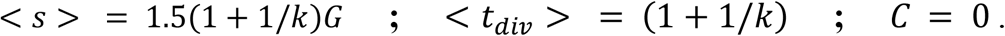

In the slow-growth limit with growth rate *G*, cells are almost exclusively below the size-threshold, and hence have division time set by the mean length of G1, plus the length of S/G2/M, plus the time it takes to grow up to the size threshold. Solving for the division time yields the following expressions:

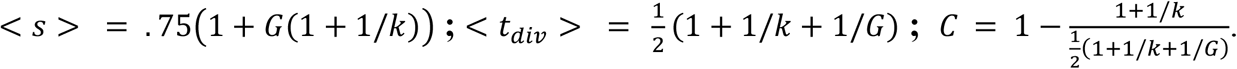

These expressions are verified against numerical results (SI Fig. S7).

### Numerical Simulation of G1-Sizer Model

In order to further understand the implications of the phenomenological G1-sizer model in a tissue context, we implemented a stochastic agent-based simulation of growing cell monolayers. In this simulation, each cell carries an index i, as well as two quantities a_i_ and p_i_, representing the size and cell cycle phase respectively. Time and size are in non-dimensional units during the simulation and are converted to dimensional units for analysis after the simulation. We do this by associating 1 unit of simulation time to be the length of a typical S phase (10 hours), and 1 unit of simulation size to be the volume at which it is seen experimentally that cells transition from a size-dependent to a size-independent division rate (1250 um^3^). We run our simulations with constant timesteps dt of .005. In the specific use case of Fig. 5E, we verified that the results obtained with a timestep of .0005 were not quantitatively different from those generated at dt = .005.

At every step in the simulation, growth of cells is advanced by changing each a_i_ by an amount *G* *dt, where *G* is the growth rate of the cell. *G* is in principle a function of the parameters of the cell itself, such as a_i_ and p_i_, as well as global parameters such as the total number of cells N and the total area of cells A. We observed similar results in a model with exponential single-cell growth (SI Fig. S8).

Division of cells is regulated by a size-dependent probability of entrance into S-phase. If a cell exceeds an area of 1, it enters S at a rate of 3, at which point its phase variable p_i_ is set to a value of 1/dt. At every step in the simulation, the phase variable p_i_ decreases by 1, and once p_i_ reaches 0, the cell’s S/G2/M phase is completed. Therefore, following stochastic initiation, each cell experiences S/G2/M phase as being a deterministic time of 1. We choose a G1 entrance rate of 3 so that the average total time of the cell-cycle in the size-independent regime is matched between simulation and experiment. At the point that a cell exits M, the cell’s size a_i_ is reduced by a factor of 2, and an additional new cell is created with size a_i_/2.

These two elements – size-dependent S entrance, and growth of cells – constitute the core of how we advance single-cell trajectories through time. We explore the implications of this framework in two categories of simulations – ensemble simulations, and single-cell simulations. In single-cell simulations, we track the trajectory of only one cell, following only one daughter cell after division. In ensemble simulations, we track a whole population of cells, and the growth rate can therefore depend on quantities like N, the total number of cells in the population.

In Fig. 5D, we show example single-cell trajectories with two different constant growth rates *G*. To exhibit timer-like behavior, we simulated cells with a growth rate g = 1.6. To demonstrate sizer-like behavior, we simulated cells with a growth rate *G* = 0.1. In this particular set of simulations, the S entrance rate was set to 10, to more clearly demonstrate the differences between the two growth regimes. Cells are initialized with a size uniformly drawn from 1 to 2, and a phase p_i_ that is either 50% uniformly distributed between 0 and 1/dt, or 50% p_i_ = -1. Trajectories are simulated for 50 units of simulation time, though only a small fraction is shown of those trajectories.

In Fig. 5E, we use single-cell simulations to look at the relation between cell size immediately post-division versus immediately pre-division in the subsequent round as a function of growth rate. We did this for 4 growth rates *G* = 1.0, 0.7, 0.2, and 0.05. Each growth rate was simulated with 400 simulation replicates, each run for 40 units of simulation time. We initialize each simulation with p_i_ = -1, and a random size uniformly distributed between 1 and 2, and allow cells to grow with *g* = 1 for 10 units of simulation time before switching to the simulation specific growth rate. After another 10 units of simulation time, we begin recording the size of a cell immediately post-division, and immediately pre-division. Confinement values are estimated as

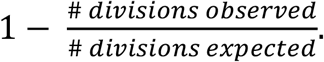

In Fig. 5F, we show results from an ensemble simulation of cells. Ensembles are initialized from 4 cells with sizes uniformly drawn from 1 to 2, and then normalized such that they have total area 8. Cell phases are either 20% uniformly distributed between 0 and 1/dt, or 80% p_i_ = -1. We first allow the cells to expand in an unconstrained way, i.e. we grow each cell in our simulation with a growth rate *G*, which is drawn for each cell and each time point from a uniform distribution with support between .9 and 1.1. When the ensemble of cells collectively exceeds a critical total size, we quench all their growth rates to 0, and therefore only division occurs from beyond that time point, which we set to be t = 0. We perform these simulations for three different critical total sizes 500, 1200, and 4700, with 20, 10, and 5 simulation replicates respectively. After the critical size is reached, we can track the ensemble distribution of areas as a function of time, which we do for 4 units of simulation time, corresponding to 48 hours of real time. We did not notice any significant differences in the distributions as a function of the total size at which we quench the growth rate.

## Notes

### Competing Interest Statement

The authors have declared no competing interest.

### Summary of Updates

Methods section added

