## Supplemental figures for "Tissue confinement regulates cell growth and size in epithelia"

This PDF file includes:
Figures S1 to S11 p2-12

Other supplementary materials for this manuscript include the following: Table S1

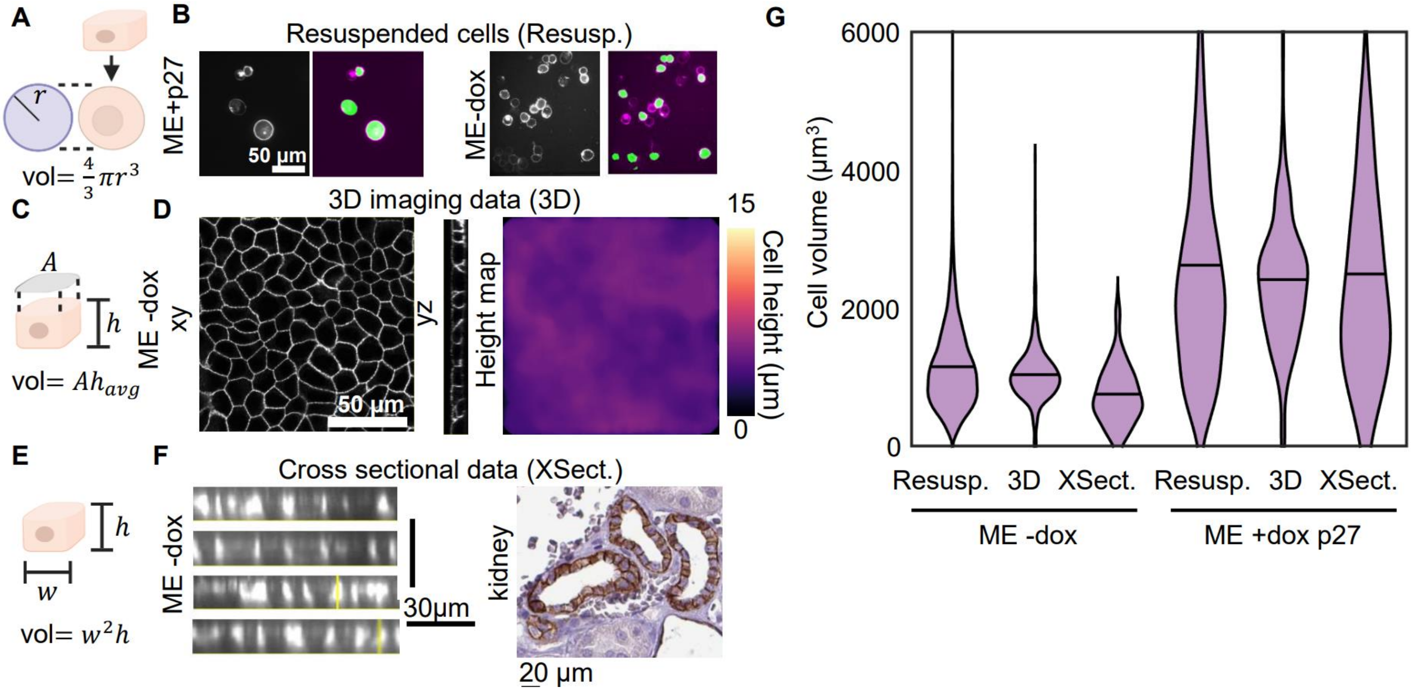


**Figure S1: Different volume measurement methods give consistent results**

We used several methods to estimate the cell volume across different experiments. While the measurement in 3D is the most accurate method of measuring the cell volume it is not feasible in all experiments. We verified that our other methods of approximation in resuspension and cross section yield the same results as full 3D imaging. (A) Schematic of cell volume measurement in resuspension (Resusp.). The resuspended cell is nearly spherical, and a measurement of the radius is used to determine the cell volume (B) Labeled membranes of resuspendend MDCK cells and corresponding cell segmentation. (C) Schematic of cell volume measurements in 3D – a height map is constructed by measuring the intensity profile at each pixel to identify the top and bottom membrane. The average cell height is multiplied by the cell cross sectional area to obtain the volume (D) 3D z-stack of labeled MDCK cell membranes in XY and YZ views. A height map measured from membranes in the image is displayed and used to measure volume from 2D segmented cell area. (E) schematic cell volume measurements in cross section (XSect.) – cells perpendicular to the imaging plane are identified and the length and width of the cell area measured to estimate the cell volume (F) images of MDCK cell membranes in cross sectional and kidney histology from Human Protein Atlas (image credit: Human Protein Atlas) stained for E-cadherin (CDH1) show cross sectional views of human kidney cells (G) quantification of cell volume using each method for MDCK tet-on P27 cells in ME-dox and ME+dox conditions. ($N_{R-}$=12641(5), $N_{3D-}$=1405(2), $N_{Xsect-}$=78(1), $N_{R+}$=1136(3), $N_{3D+}$=,324(2) $N_{Xsect+}$=59(1))

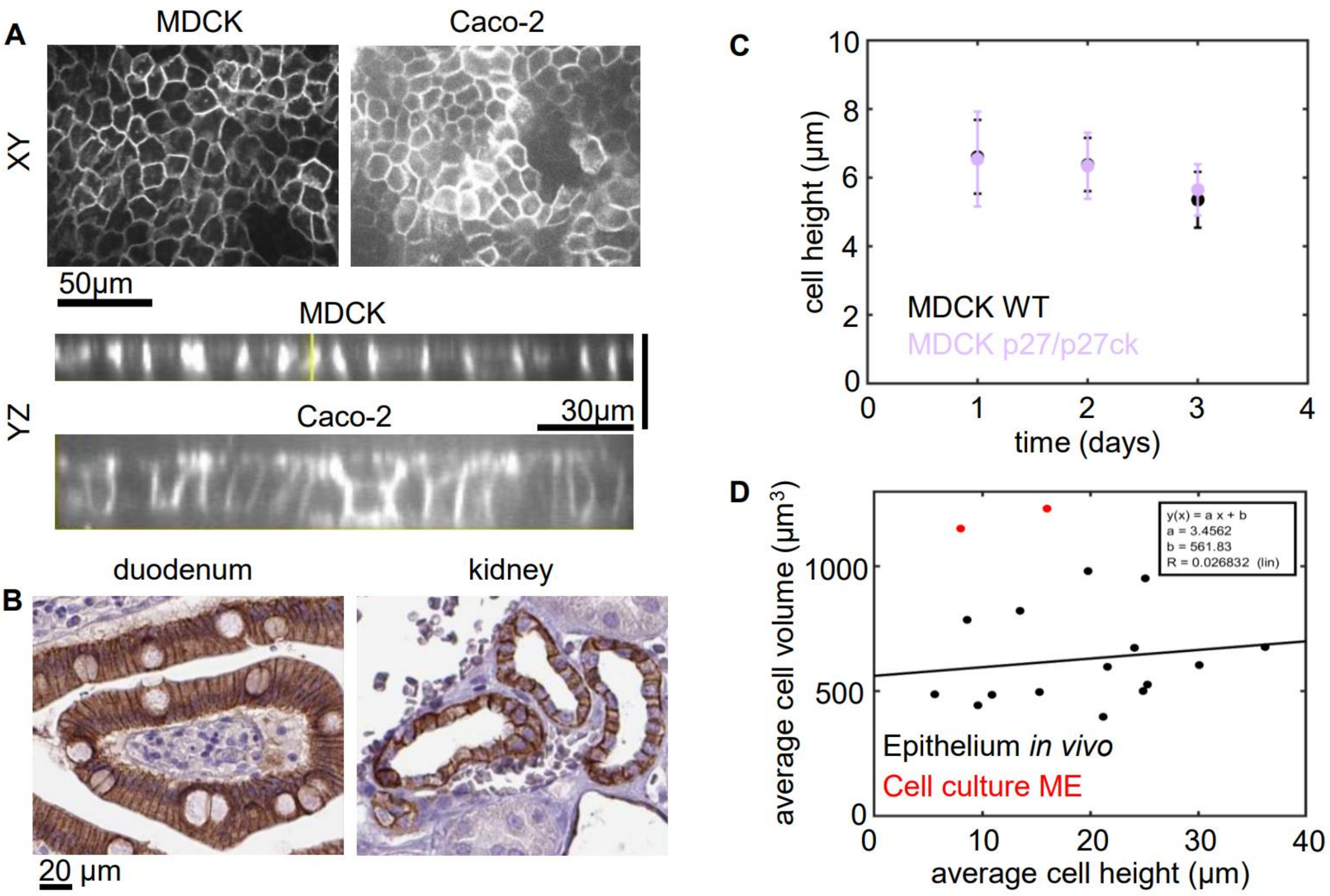


**Figure S2: Cell height is consistent for a given cell type but varies between tissues**
Across our dataset we observed a large variation in cell height. We verified that across experiments and different time points with the same cell line the height remains nearly consistent. We also show that the difference in volume between different cell types is not a consequence of differences in height. (A) MDCK and Caco-2 cell membranes showing XY and YZ views of monolayers. (B) Histology section images from Human Protein Atlas (image credit: Human Protein Atlas) of Duodenum and Kidney tissue stained for E-cadherin (CDH1). (C) Cell height of MDCK monolayers across different time points. Data are from 3D segmentation. Error bar shows standard deviation of 20 different fields of view containing >100 cells from 1 experiment each. P27/p27ck are monolayers made from a mixture of Tet-On p27 and Tet-On p27ck cells with doxycycline added at t=0. (D) Plot of cell height vs volume measured in cross section across 15 different tissues *in vivo* analyzed in Fig. 1B and two cell lines cultured as ME *in vitro* (MDCK and CACO-2). Black line shows linear fit to *in vivo* data. Variations in cell height do not explain the variation in cell volume.

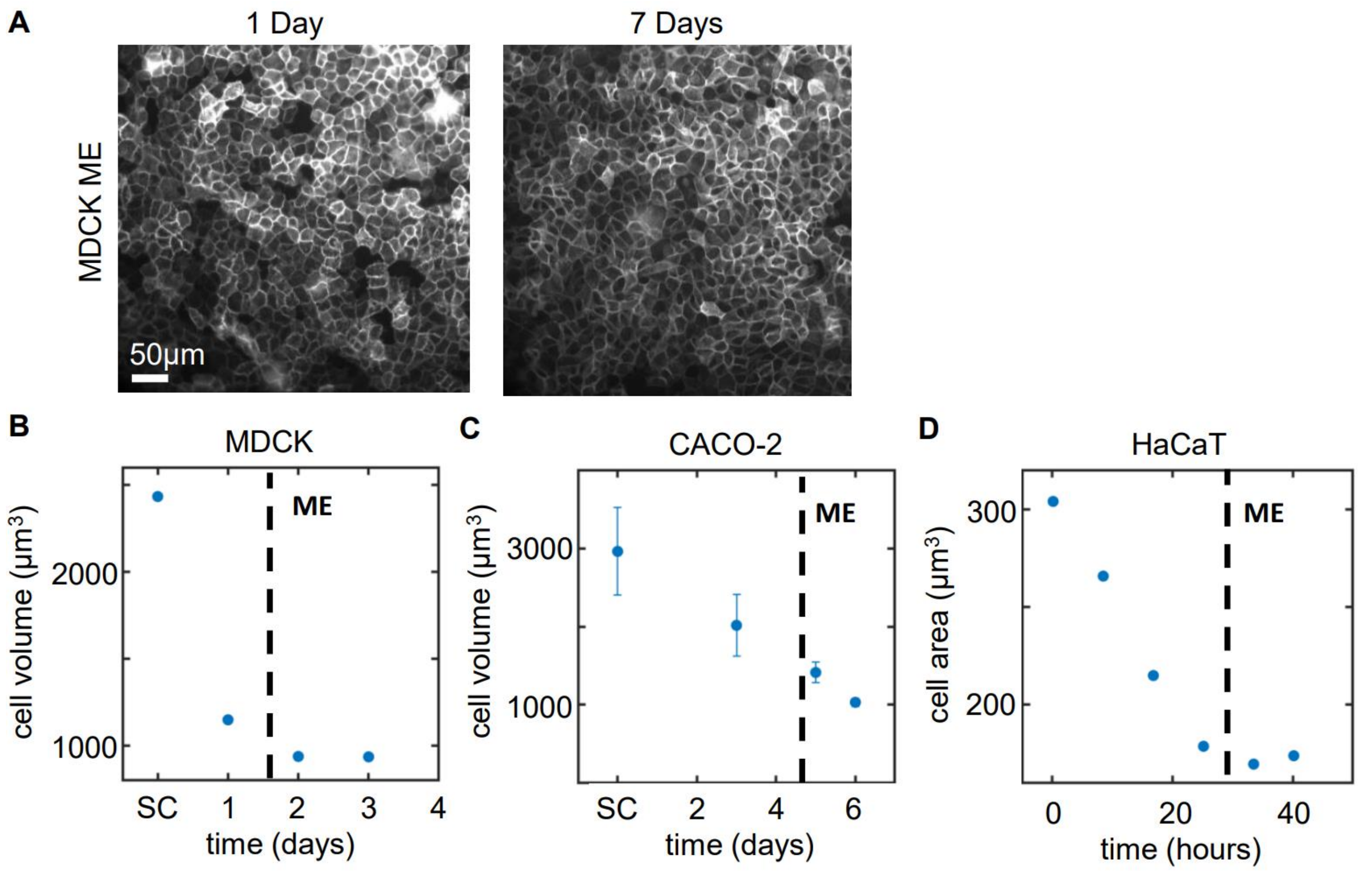


**Figure S3: Cell volume reaches a plateau in mature epithelium (ME)**
We wanted to confirm that in the mature epithelium there are no changes in cell volume. We performed volume measurements across different time points for each cell line and define what time points we consider the system to reach ME by for each cell line. Due to differences in division rates this varies but all cell lines show a decreasing size over time leading to a plateau. (A) MDCK cells with labeled membranes in monolayers imaged at ME+1 day and ME+7 days (B) Measurements of MDCK cell volume at different time points in. Data are from 3D segmentation of >200 cells at each time point from >10 fields of view in 1 experiment (C) CACO-2 cell volume measured in resuspension at different time points. Error bar shows standard deviation from 2 experimental replicates. (D) Measurement of HaCaT cell area at different time points. Each point is the average area from n=50 cells from 3 different fields of view in a single time lapse experiment. In all experiments t=0 is the onset of confluence experiments and the point of ME is indicated with a dashed line.

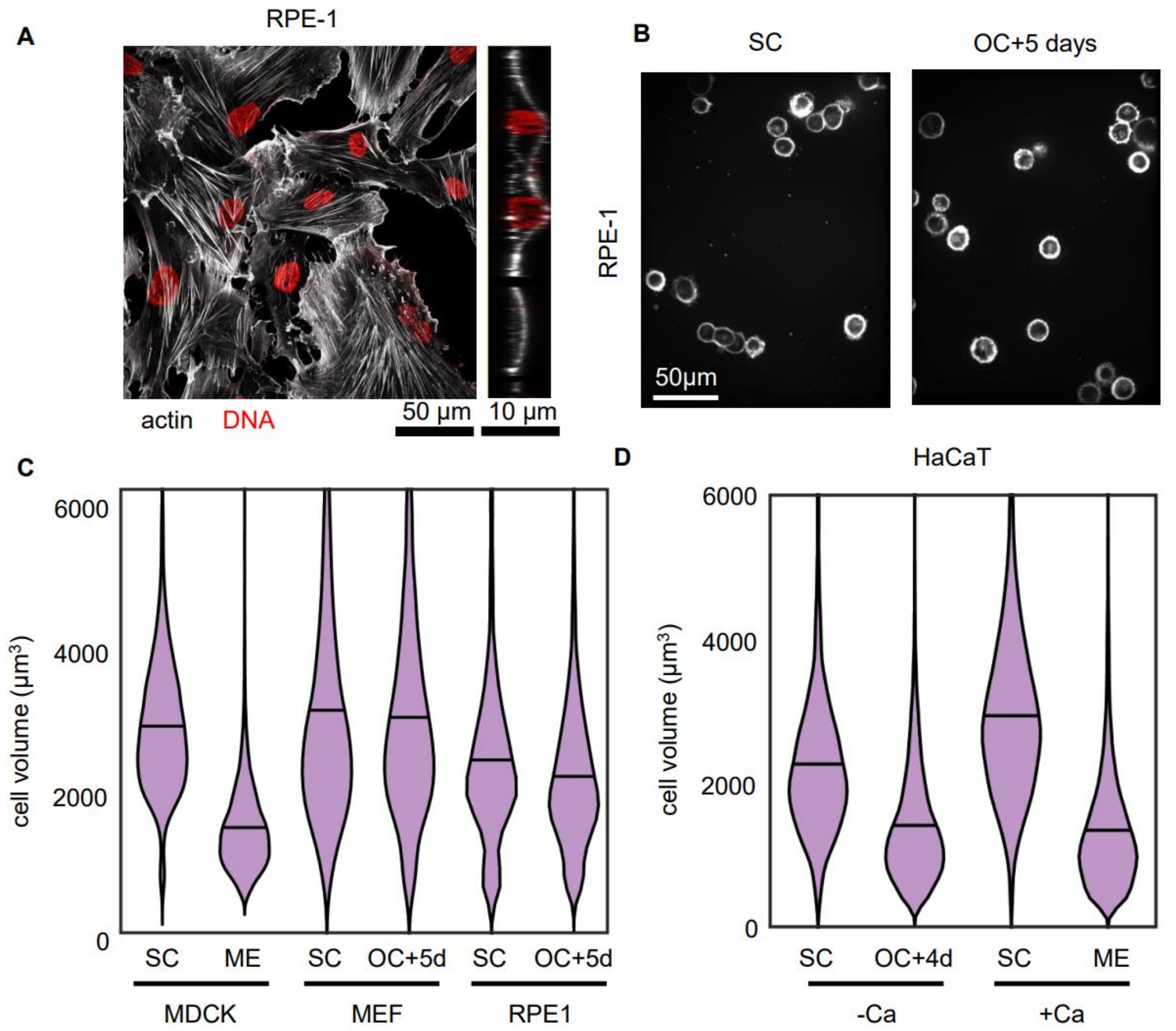


**Figure S4: Cell volume does not change during contact inhibition of non-epithelial cells**
We wanted to determine if the uncoupling of cell growth and division also extends to other cell types which would not form epithelium *in vivo*. We tested two cell lines with fibroblast-like morphology and found that these do not show cell size changes. HaCaT cells are routinely cultured in the absence of calcium which inhibits junction maturation and differentiation, but this did not inhibit cell size reduction in HaCaT cell under confinement. Future work is required to understand this difference in behavior for epithelial and non-epithelial cells. (A) staining for actin and DNA in retinal pigment epithelial (RPE-1) cells. Cells do not form coherent colonies or show epithelial morphology (B) RPE-1 cells in suspension labeled with CellMask Deep Red from 5 day old cultures plated at low density to remain SC all 5 days or high density to reach OC at day 0. (C) Volume measured in resuspension of MDCK, RPE-1 and mouse embryonic fibroblast (MEF) cells SC and ME-like culture conditions. SC – subconfluent, ME- mature epithelium at day 4, OC+5d- 5 days after the onset of confluence N = #cells (#exp) ($N_{MDCK-SC}$=4940(3), $N_{MDCK-ME}$=12641(5), $N_{MEF-SC}$=1459(1), $N_{MEF-OC5}$=1352(1), $N_{RPE-SC}$=906(2), $N_{RPE-OC5}$=3429(2)) (D) Volume measured in resuspension of HaCaT cells in SC and ME cultures with and without calcium. -Ca – 40uM calcium, +Ca – 1.8mM calcium ($N_{SC-Ca}$=3642(2), $N_{OC4-Ca}$=23996(2), $N_{SC+Ca}$=9248(2), $N_{ME+Ca}$=13451(2))

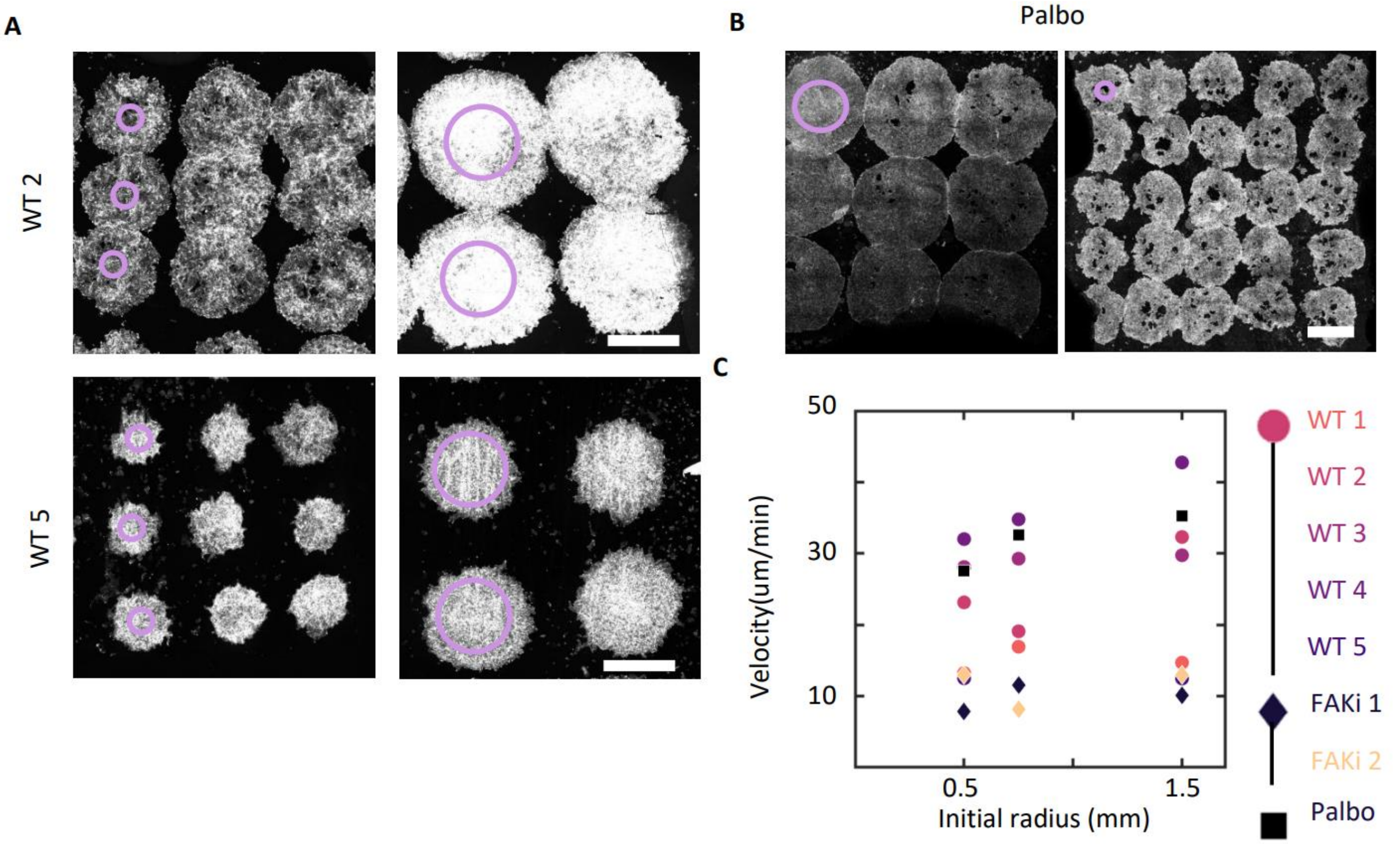


**Figure S5: Colony expansion rate varies between experiments and is not cell division dependent** Between different expanding monolayers we observed different migration velocities. We characterized the variation in speed across all experiments. We also tested that the expansion of colonies is not driven by cell division by inhibiting the cell cycle with Palbociclib (A) Expanding colonies labeled with CellTrace Far Red with initial radius 0.5mm and 1.5mm in two different experimental replicates. Migration velocity is consistent within the same experiment but varies between experiments (B) Expanding colonies labeled with CellMask Deep Red treated with 1µM Palbociclib to block cell division. Migration rate is ~30µm/hr equal to the highest rate observed in control conditions. (C) Colony expansion speeds in each experiment for different initial sizes. Expansion occurs for 48 hours and the difference between initial and final radii is used to calculate the velocity. +FAKi is 500nM PND1186. Palbo is 1µM Palbociclib

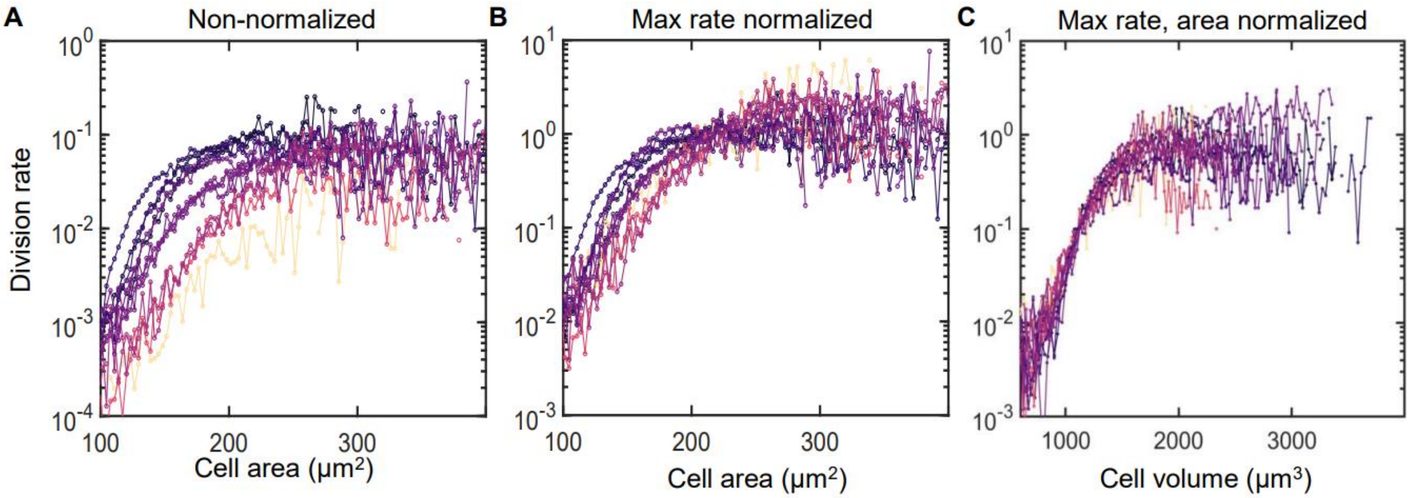


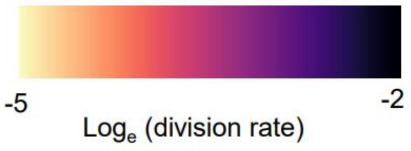


| Expt# | Cell type | substrate | inhibitor | Division rate |
|---|---|---|---|---|
| 1 | MDCK | 2mg/ml col gel | | 0.094 |
| 2 | MDCK | 2mg/ml col gel | | 0.088 |
| 3 | MDCK | 2mg/ml col gel | | 0.046 |
| 4 | MDCK | 2mg/ml col gel | | 0.067 |
| 5 | MDCK | 2mg/ml cross linked col gel | | 0.078 |
| 6 | MDCK | 4mg/ml cross linked col gel | | 0.047 |
| 7 | MDCK | Glass | | 0.033 |
| 8 | MDCK | 2mg/ml col gel | 25μM rac inh. | 0.048 |
| 9 | MDCK | 2mg/ml col gel | 50μM rac inh. | 0.030 |
| 10 | MDCK | 2mg/ml col gel | 500nM FAK inh. | 0.022 |
| 11 | MDCK | 2mg/ml col gel | 3um $Mn^{2+}$ | 0.046 |
| 12 | CACO-2 | 2mg/ml col gel | | 0.008 |

**Figure S6: Cell division rate shows a similar size dependence across a range of experimental conditions**

We reanalyzed a previously published dataset to see how cell division rates vary with cell size in epithelium. A subset of the data is presented in Fig. 5A but complete set of data analyzed is presented in this figure. We see a consistent relationship between division rate and area despite differences in the overall division rate across experiments. (A-C) Plots of division rates against cell size across 12 timelapse imaging experiments that are previously published by Devany et. al. 2021. Plots show data from 4 replicates of MDCK cells on 2mg/ml collagen gels, data with variation in substrate stiffness, with inhibitors of RAC1 (NSC 23766) and FAK (PND1106), and with CACO-2 cells. See table for full experimental details. Data are plotted non-normalized in A, normalized by the maximum rate in B, and normalized by both maximum rate and the area is normalized so the rate 0.1 is at a volume of 1200 μm³. Curves are colored according to the maximum division rate. Each division rate curve is generated from tracking >300 division events from each experiment over a timelapse imaging series of at least 16 hours (see Methods).

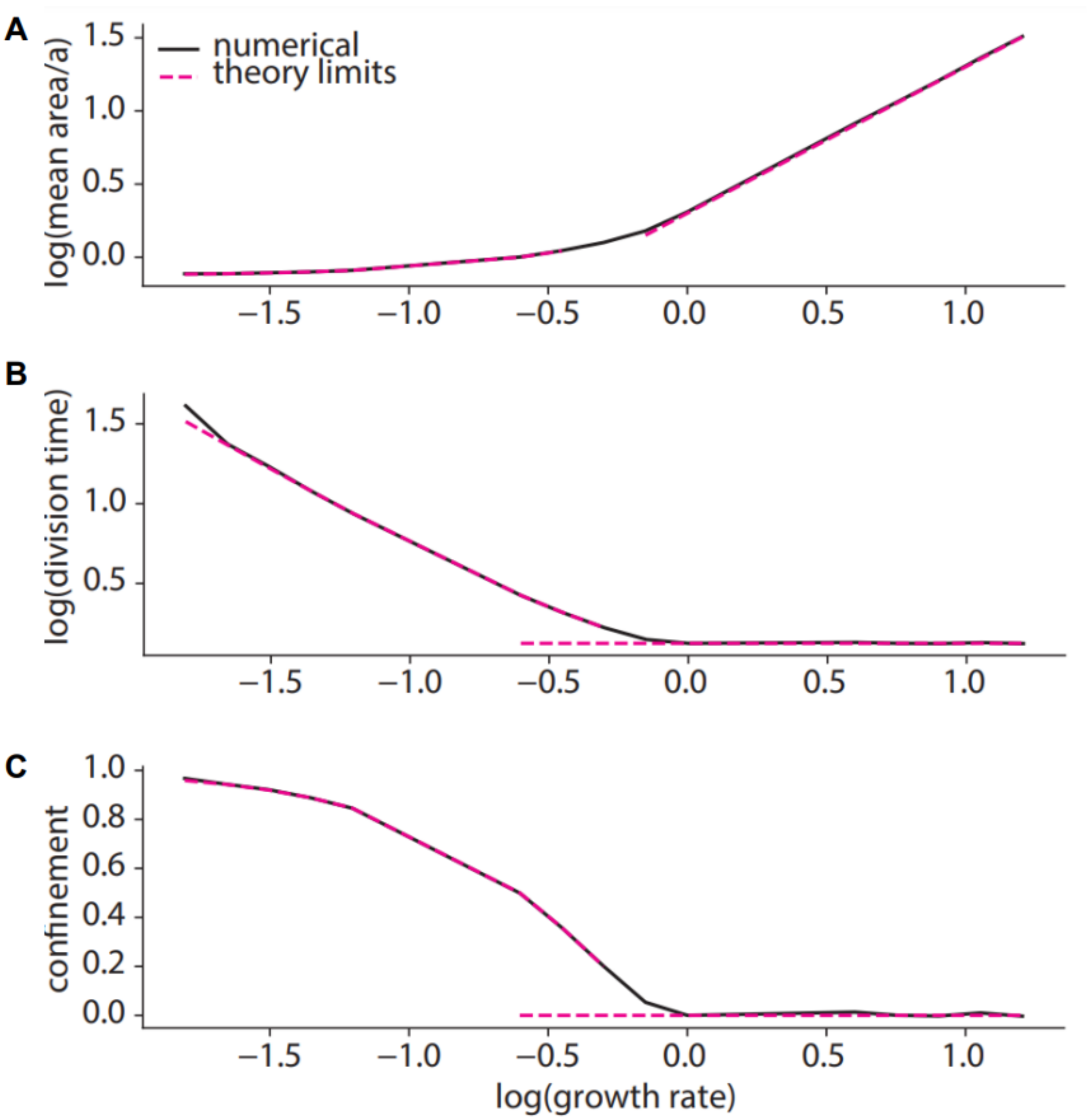


**Figure S7: G1-sizer control generates timer and sizer behaviors depending on growth rate**
We numerically simulate single-cell trajectories (see Methods) as a function of growth rate. We compare our numerical results to analytical expressions (see Methods) derived in the fast- and slow-growth limits of the model. (A) Numerical simulation of mean non-dimensional cell size (black) reveals two distinct regimes: a sizer regime where cell size is only weakly dependent on growth, and a timer regime where size is linear with growth. This agrees well with theoretical expectations in the limits of fast and slow growth (pink). (B) Numerical simulation of mean division time (black) reveals two distinct regimes: a sizer regime where division time diverges with decreasing growth, and a timer regime where division time is independent of growth. This agrees well with theoretical expectations in the limits of fast and slow growth (pink). (C) Numerical simulations (black) reveal a confinement transition when cells switch from sizer to timer behavior: confinement is non-zero for slow-growth sizers, and zero for fast-growth timers. This agrees well with theoretical expectations in the limits of fast and slow growth (pink).

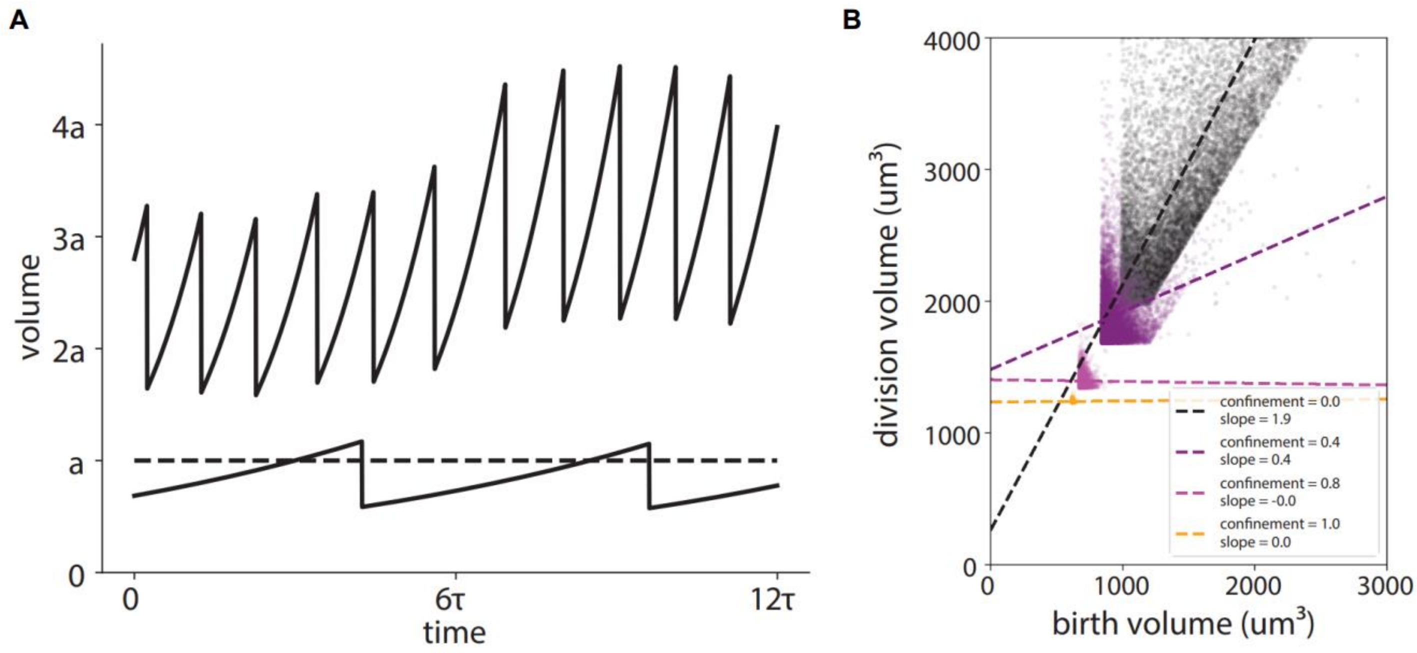


**Figure S8: A G1-sizer with single-cell exponential growth shows a transition between timer and sizer behavior**

To verify that the transition between timer and sizer behavior in our model is not an artifact of linear cell growth, we simulate a model with exponential growth, where cell size $a_i$ increases by an amount $a_i * G * dt$ at each time step. (A) Numerical simulation of cells with $G = .74$ (upper solid line) and $g = .14$ (lower solid line) reveals qualitatively distinct timer and sizer behaviors as a function of growth relative to the G1-exit size cutoff (dashed line). The $G = .74$ trajectory displays the increasing variance of cell size characteristic of timer control with exponential single-cell growth. (B) Plotting numerically simulated cell birth sizes vs division sizes reveals a transition from timer to sizer behavior as a function of confinement. The slope of the linear best fit lines (dotted) to the data goes from timer-like (~2) to sizer-like (~0) as confinement increases. Growth rates $G = \{ 0.5, 0.33, 0.11, 0.025 \}$ going from small to large confinement.

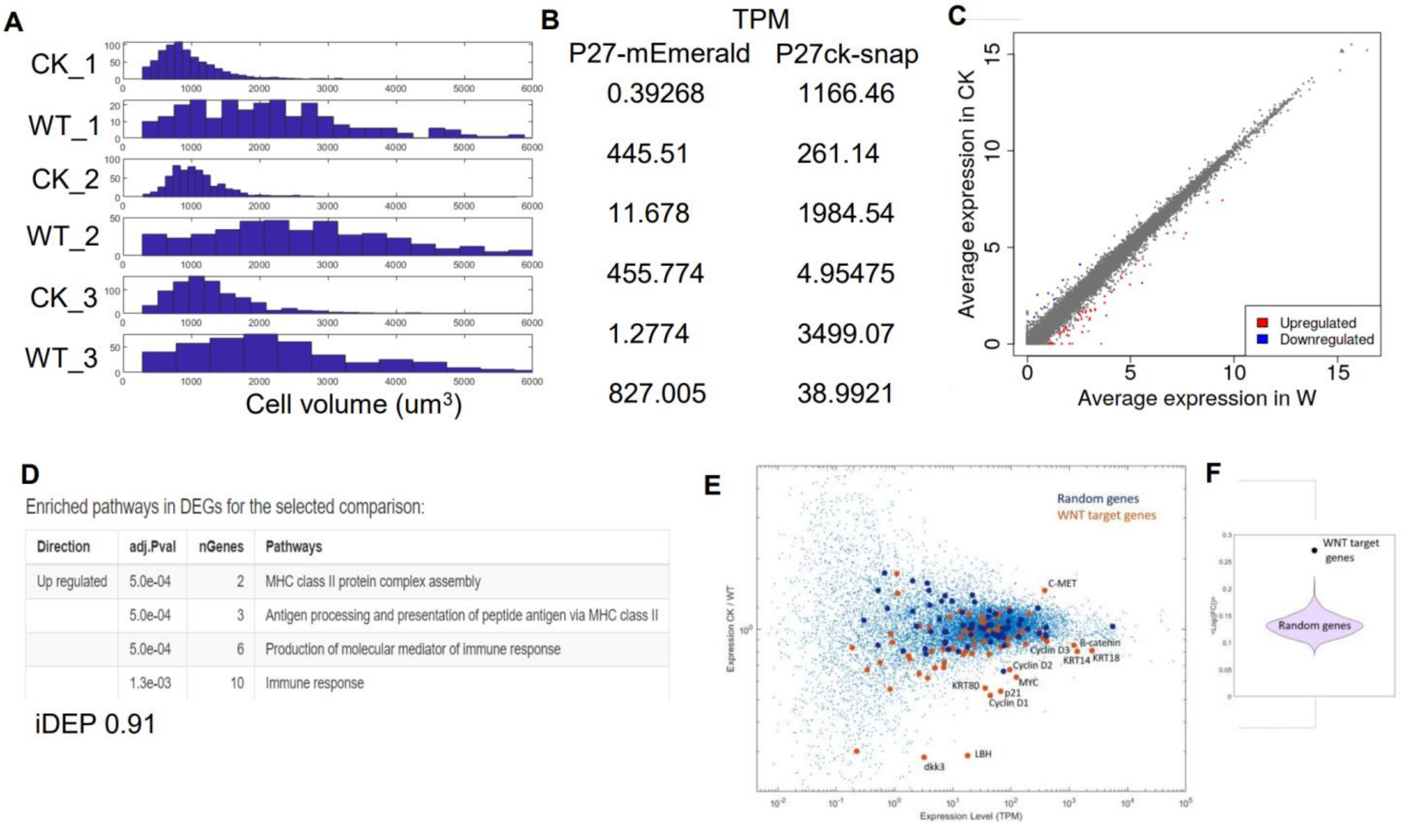


**Figure S9: RNA expression is similar between large and small cells but shows weak immune and WNT signature**

We performed RNA sequencing on monolayers with different cell size determined by inhibiting cell division. The data is presented in Fig 6B. This figure includes additional analysis performed on the data which identified some weak transcriptional signatures that we do not interpret to explain the size dependent cell cycle arrest (A) Distributions of cell volumes measured in suspension from each RNA seq experiment (CK – Tet-On Snaptag-p27ck cells +dox, WT – Tet-On GFP-p27 cell +dox). Dox was added at t=0 and RNA was collected 5 days later at ME+4 days. 3 samples were made for each experiment and 2 were pooled for RNA extraction and sequencing while the other sample was used to measure the volume distribution (B) expression level of GFP-p27 and Snaptag-p27ck found in each of the samples. Each data set is one experimental replicate with 2 technical replicates. (C) Plot of RNA expression from Fig. 6b with genes which are significantly upregulated or downregulated ($P<0.05$) highlighted. Data are averaged from 3 experimental replicates each with 2 technical replicates (D) Pathway enrichment analysis of genes from RNA sequencing data showing all pathways with significant upregulation or downregulation. Analysis performed using iDEP 0.91 (see Methods) (E) plot showing the ratio of gene expression vs the average expression. 65 WNT target genes from The WNT Homepage are plotted in orange while a random set of 65 genes is shown in blue (F) violin plot of average fold change in 1000 sets of 65 random genes compared to the average fold change of WNT target genes in E.

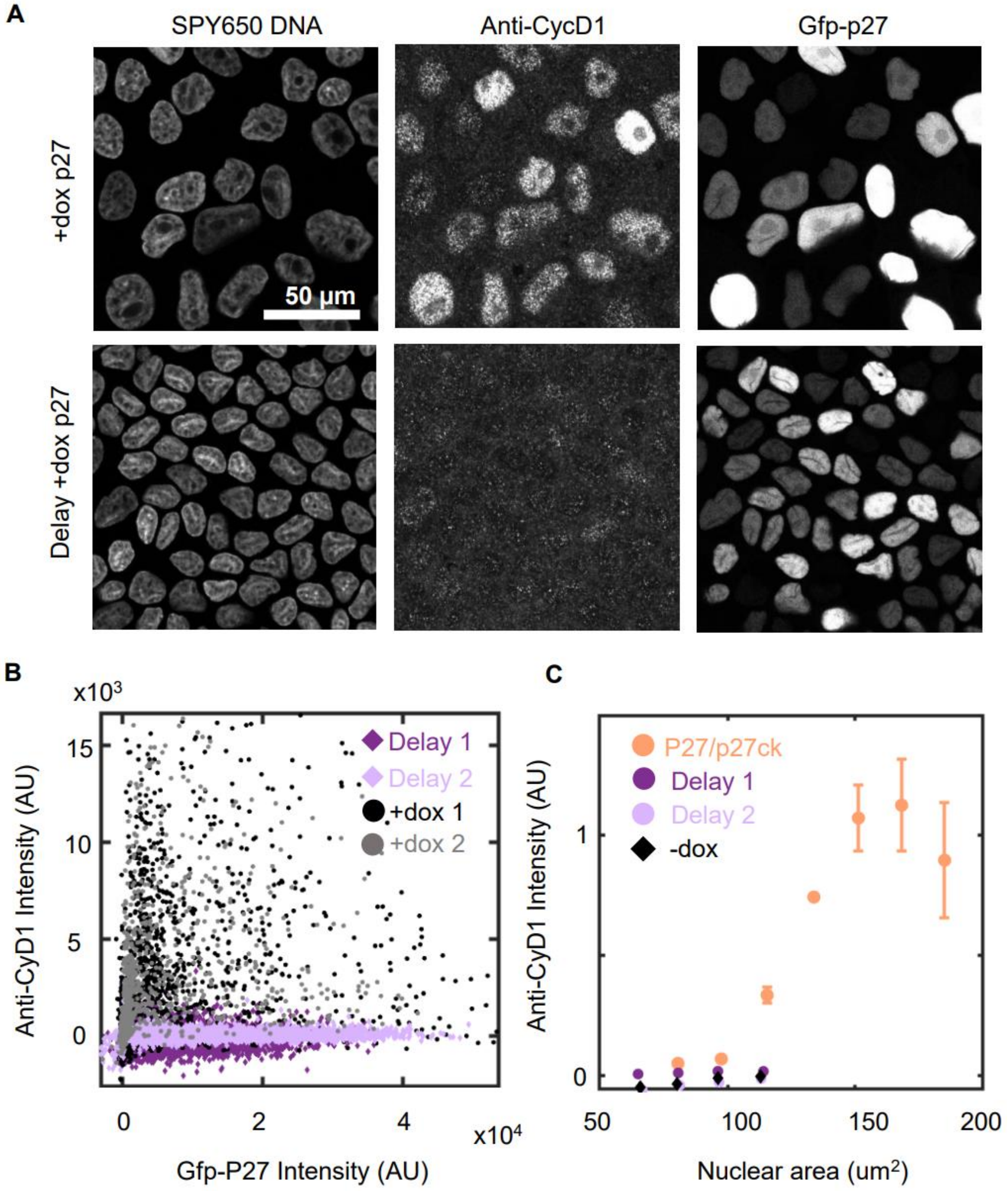


**Figure S10: Cyclin D1 level is independent of p27 expression**

Due to a direct binding interaction between p27 and Cyclin D we were concerned that overexpression of p27 may alter the levels of Cyclin D1 found in normal cells. We performed controls where we manipulate cell size independent of p27 levels and show that there is no resulting effect on expression levels of Cyclin D1 (A) MDCK Tet-On GFP-p27 monolayers labeled with spy650 DNA, anti-cyclin D1 and GFP-p27 under conditions where doxycycline is added at day 0 (+dox p27) or at day 2 (Delay +dox p27) (B) plot of anti-Cyclin D1 nuclear intensity vs GFP-p27 nuclear intensity in +dox P27 or delay +dox P27 conditions (C) Plot of anti-Cyclin D1 nuclear intensity in p27/p27ck coculture, delay +dox p27 and -dox condition. Delay+dox p27 and -dox show similar behavior suggesting that p27 expression does not affect the relationship between Cyclin D1 levels and cell size ($N_{+dox1}$ =1820(1) $N_{+dox2}$ = 1112(1) $N_{Delay1}$ = 3479(1), $N_{Delay2}$ =4036(1), $N_{-dox}$ = 4155(1) $N_{p27/p27ck}$= 11080(4))

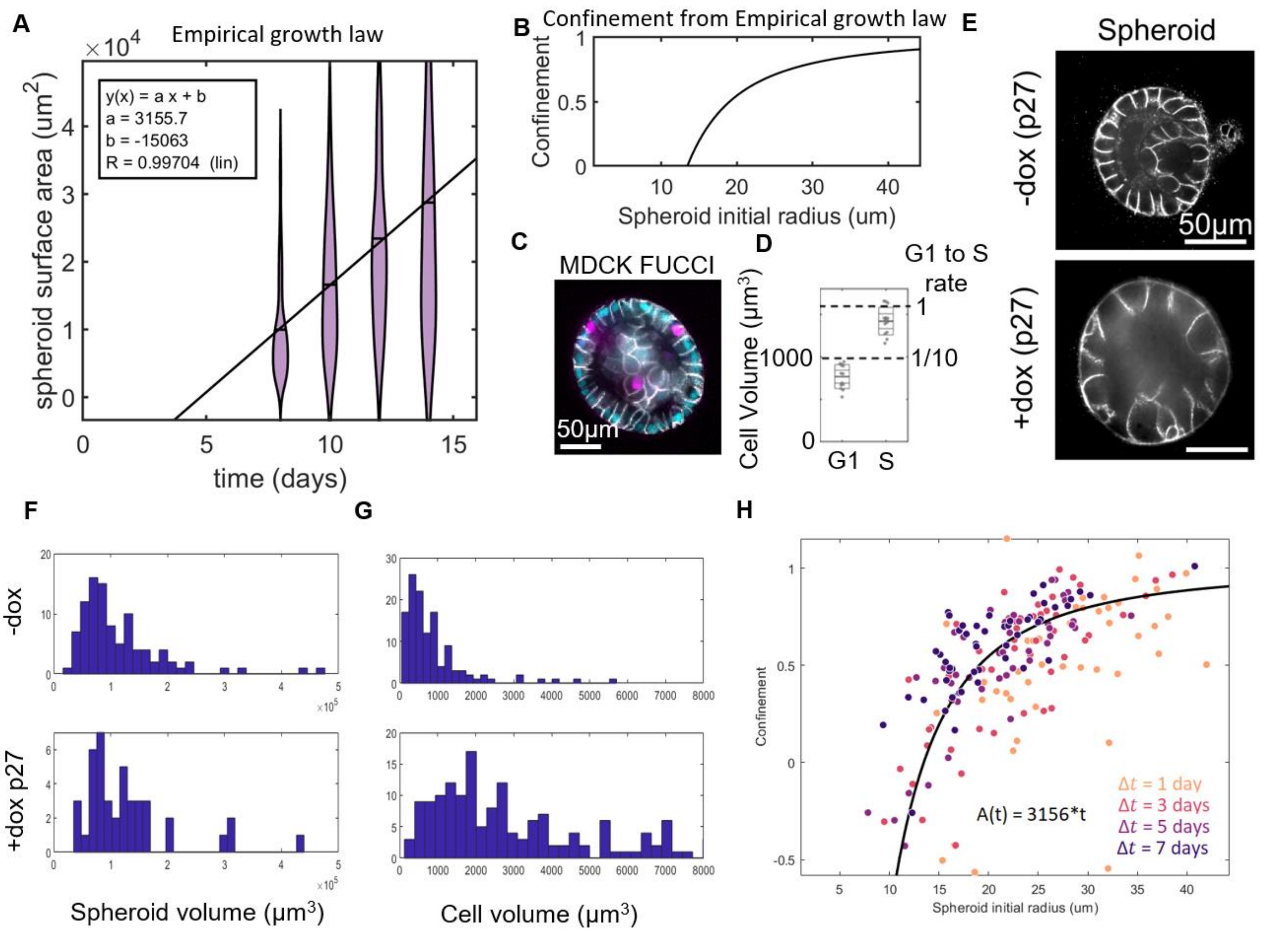


**Figure S11: Epithelial spheroids are amenable to confinement analysis and show high confinement** We measured the growth rate of MDCK spheroids and of cells within spheroids to demonstrate another application of our confinement framework. In this context the spheroid expansion is not driven by migration as it is in the expanding monolayers and the growth law is not known. By measuring the behavior of spheroids, we find an empirical growth law and use this to predict the confinement. This confinement agrees with an independent growth measurements on single cells. (A) Distribution of epithelial spheroid size measured at different time points. Linear fit show the surface area of the spheroid increases with time (B) Confinement as a function of size calculated using the linear growth of spheroid area empirically determined from A (C) Epithelial spheroid of MDCK FUCCI cells with labeled cell membranes (D) Quantification of cell volume of randomly selected cells in G1 and S phase in MDCK spheroids at day 8 (N=13 cells from 1 experiment in each condition). Dotted lines show volumes where the division rate reaches is maximal or 1/10 the maximum from Fig. 5A (E) Representative images of spheroids formed with Tet-On p27 MDCK cells without dox (-dox) or with dox added at t = 5 days. Spheroids were imaged at t = 10 days (F) spheroid volumes in -dox and +dox p27 conditions. Data are from >40 spheroids each in 1 experiment (G) distribution of cell areas in -dox and +dox p27 conditions. Data are from >100 cells each in 1 experiment (F) Measured confinement for individual spheroids from the cell growth averaged across a spheroid measured from changes in cell size under p27 induction compared with the growth model shown in B (see Methods). Cell size in each spheroid is measured after $\Delta t$ in the presence of dox and compared with the expected size in -dox conditions ~1000μm$^3$. The difference in cell size divided by $\Delta t$ gives the cell growth rate which is compared to the single cell rate of MDCK to give the confinement.
